# Exploiting selection at linked sites to infer the rate and strength of adaptation

**DOI:** 10.1101/427633

**Authors:** Lawrence H. Uricchio, Dmitri A. Petrov, David Enard

**Affiliations:** Department of Biology, Stanford University, Stanford, CA 94305; Department of Ecology and Evolutionary Biology, University of Arizona, Tucson, AZ 85721

## Abstract

Genomic data encodes past evolutionary events and has the potential to reveal the strength, rate, and biological drivers of adaptation. However, robust estimation of adaptation rate (*α*) and adaptation strength remains a challenging problem because evolutionary processes such as demography, linkage, and non-neutral polymorphism can confound inference. Here, we exploit the influence of background selection to reduce the fixation rate of weakly-beneficial alleles to jointly infer the strength and rate of adaptation. We develop a novel MK-based method (ABC-MK) to infer adaptation rate and strength, and estimate *α* = 0.135 in human protein-coding sequences, 72% of which is contributed by weakly adaptive variants. We show that in this adaptation regime *α* is reduced ≈ 25% by linkage genome-wide. Moreover, we show that virus-interacting proteins (VIPs) undergo adaptation that is both stronger and nearly twice as frequent as the genome average (*α* = 0.224, 56% due to strongly-beneficial alleles). Our results suggest that while most adaptation in human proteins is weakly-beneficial, adaptation to viruses is often strongly-beneficial. Our method provides a robust framework for estimating adaptation rate and strength across species.

## Introduction

The relative importance of selection and drift in driving species’ diversification has been a matter of debate since the origins of evolutionary biology. In Darwin and Wallace’s formulations of evolutionary theory, natural selection is the predominant driver of the accumulation of differences between species (*1,2*). Subsequent theorists argued that random genetic drift could be a more important contributor to differences between species (*3–6*), with chance differences accumulated over time due to reproductive isolation between populations. Although it is now clear that natural selection plays a substantial role in both diversification and constraint in many species (*7–10*), considerable uncertainty remains about the relative importance of stochastic drift, mutation, selection, and linkage, with no clear consensus among evolutionary geneticists (*11–15*). A better mechanistic understanding of these processes and how they jointly shape genetic diversity could help to resolve old evolutionary puzzles, such as the narrow range of observed genetic diversity across species (*16*) and the apparently low rate of adaptation in primates (*17*).

With the exception of rapidly evolving microbial species, most adaptation events occur too slowly to be directly observed over the timescale of a scientific study. Therefore, detailed study of the molecular basis of adaptation has required the development of computational methods to infer adaptation rate (denoted *α*, defined as the proportion of fixed differences between species that confer fitness benefits) directly from genetic sequence data. Most existing approaches derive from the McDonald-Kreitman (MK) test (*7,18*) and related Poisson random field framework (*19*), both of which use divergence and polymorphism data to infer adaptation rates. Note that a recent approach uses polymorphism data alone to infer the distribution of fitness effects of fixing mutations (*20*). The critical idea behind each of these methods is to compare evidence for differentiation at alleles that are likely to have fitness effects (e.g., nonsynonymous alleles that change protein function by altering the amino acid sequence) to alleles that are less likely to have fitness effects (e.g., synonymous alleles that do not change the amino acid sequence of proteins).

In the classic MK framework, the rate of divergence at putatively functional sites (*D_N_*, often defined as nonsynonymous differences within proteins) is compared to putatively neutral diverged sites (*D_S_*, often defined as synonymous differences). Polymorphic sites within both the functional and non-functional class (*P_N_* and *P_S_*, respectively) are used as a background to calibrate the expected rate of divergence under a neutral model. If mutations at functional sites are assumed to be either virtually lethal or neutral, then the ratio 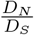 has the same expected value as 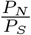 given that virtually lethal mutations contribute to neither *P_N_* nor *D_N_*. When 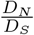 exceeds 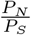, this is interpreted as evidence of adaptation because sites with functional effects on proteins are over-represented among the fixed differences relative to the neutral expectation. Smith and Eyre-Walker developed a simple equation that uses the same logic as the MK test to estimate adaptation rate *α*,

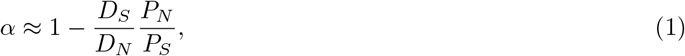

and used this approach to provide evidence for a high rate of adaptation in *Drosophila* (*18*).

Unfortunately, this elegant framework is susceptible to many biases, most notably driven by the presence of weakly deleterious polymorphism in the class *P_N_*. Deleterious polymorphism effectively makes the test overly conservative, because deleterious alleles are unlikely to ever reach fixation and therefore lead to the overestimation of the expected background rate of substitutions in the functional class. Fay *et al* introduced the idea of including only common polymorphic alleles (*e.g.*, alleles at frequency 15% or greater), which should remove many deleterious alleles (*21*) – however, this approach has been shown to provide conservative adaptation rate estimates in many contexts (*22*). More recently, Messer & Petrov showed that even removing all polymorphism below 50% is insufficient to correct this bias, especially when slightly deleterious mutations are common and the rate of adaptive evolution is high (*23*). In order to mitigate this effect, Messer & Petrov introduced the idea of the asymptotic MK test (aMK). In this implementation, 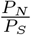 in eqn. 1 is replaced by 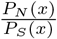, where *P_N_*(*x*) and *P_S_*(*x*) are the number of segregating nonsynonymous and synonymous alleles at frequency *x*, respectively (*23*). An exponential curve is fit to the resulting *α*(*x*) function, which can be calculated for all values of *x* in the interval (0,1) for a sample of sequenced chromosomes. The intercept of the best-fit exponential curve at *x* = 1 is a good approximation for *α*, as it effectively removes all slightly deleterious polymorphism at all frequencies. This approach was shown to be robust to both the underlying distribution of deleterious effects and recent demographic events (*23*). aMK has inspired new approaches to inferring adaptation in mitochondrial genes (*24*) and revealed a high rate of adaptation in proteins interacting with pathogens (*25*).

While aMK extends the elegant MK framework for estimating adaptation rate, it does not explicitly account for the possibility that beneficial alleles contribute to segregating polymorphism. It is unknown whether aMK is robust to the presence of weakly beneficial alleles, but there is reason to believe that beneficial alleles would be problematic because they are preferentially found at very high frequencies (*20*), and thus their effect would not be eliminated by the asymptotic procedure. The recent emphasis on adaptation from standing variation (*26–30*) and reported evidence for weakly-beneficial polymorphism in *Drosophila* (*31*) suggest that methods to infer adaptation strength over longer evolutionary time-scales are needed.

A key limitation of existing MK-based approaches is that they provide estimates of adaptation rate but not adaptation strength, and therefore it is not clear whether weakly beneficial mutations contribute substantially to the fixation process. The underlying processes driving weak and strong adaptation might differ, and the ability to separately estimate rates of weak and strong adaptation could provide insight into the biological drivers of adaptation. We hypothesized that such a method could be developed by exploiting the impact of background selection (BGS) on the fixation rate of weakly-beneficial alleles. BGS removes neutral and weakly-beneficial variation via linkage to deleterious loci (*32*), while the fixation rate of strongly-adaptive alleles is not substantially affected (*33*). Given that the strength of BGS varies widely and predictably across the human genome (*34*), a method that interrogates the rate of adaptation as a function of BGS might be able to jointly infer the rate and strength of adaptation.

Here, we probe the performance of aMK when weakly-beneficial alleles substantially contribute to segregating polymorphism, and we show that aMK underestimates *α* in this adaptation regime. We additionally show that when adaptation is weak, true *α* is predicted to vary substantially across the genome as a function of the strength of BGS. We exploit this signal of covariation between *α* and BGS in the weak-adaptation regime to develop an approximate Bayesian computation method, which we name ABC-MK, that separately infers the rate of adaptation for weakly-beneficial and strongly-beneficial alleles. Our approach and aMK rely on similar input data, but we use a model-based fitting procedure that directly accounts for BGS and weakly-beneficial alleles. We apply our method to human genetic data to provide evidence that adaptation in humans is primarily weakly-beneficial and varies as a function of BGS strength. Interestingly, adaptation rate estimates on virus-interacting proteins support a much higher rate of strong adaptation, suggesting that adaptation to viruses is both frequent and strongly fitness-increasing. We address seven potential sources of confounding, and discuss our results in light of recent research on adaptation in humans.

## Results

### *α* estimates are conservative for weakly-beneficial selection

The aMK approach is known to converge to the true *α* at high frequency under the assumption that positively selected mutations make negligible contributions to the frequency spectrum (*23*). This assumption is likely to be met when beneficial alleles confer large fitness benefits, because selective sweeps occur rapidly and beneficial alleles are rarely observed as polymorphic. However, when selection is predominantly weak, attaining a substantial *α* requires much larger mutation rates for beneficial alleles and longer average transit time to fixation, introducing the possibility that weakly-beneficial alleles will contribute non-negligibly to the frequency spectrum, even in small samples.

We tested the robustness of the aMK approach to the presence of weakly-beneficial alleles using simulation and theory. We simulated simultaneous negative and positive selection using model-based forward simulations under a range of scenarios (*35, 36*). We supposed that nonsynonymous alleles are under selection, while synonymous alleles are neutral. In each simulation, we set *α* = *α_W_* + *α_S_* = 0.2, where *α_W_* is the component of *α* due to weakly-beneficial mutations (*2Ns* = 10) and *α_S_* represents strongly-beneficial alleles (*2Ns* = 500). Note that *α* is not treated as a parameter in the analyses herein; we use analytical theory to calculate the mutation rates for deleterious alleles and advantageous alleles that result in the desired *α*, meaning that *α* is a model output and not a model input. We drew deleterious selection coefficients from a Gamma distribution inferred from human sequence data (*37*), and we varied *α_W_* from 0 to 0.2 (Fig. 1).

**Figure 1:**
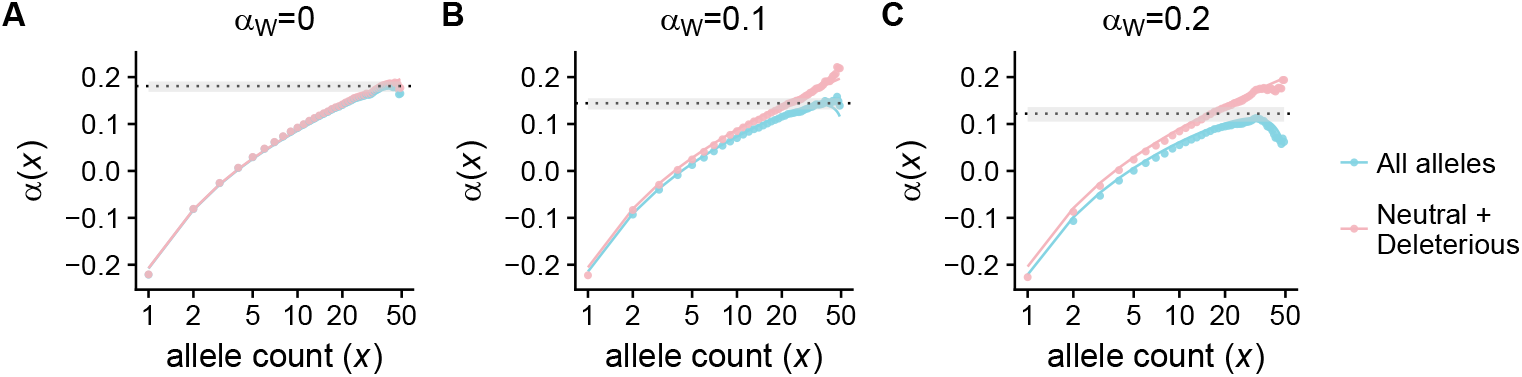
A-C: We plot *α*(*x*) as a function of allele count *x* in a sample of 50 chromosomes. The true value of *α* = 0.2 in each panel, with varying contributions from weakly (2*Ns* = 10) and strongly adaptive alleles (2*Ns* = 500). The solid lines show the results of our analytical approximation (eqn. 11), while the points show the value of *α*(*x*) from forward simulations. The blue points and curves show the calculation as applied to all polymorphic loci, while in the pink points and curves we have removed positively selected alleles from the calculation. The dotted line shows the estimated value of *α* from the simulated data using existing asymptotic-MK methods (*23, 38*), while the gray bars show the 95% confidence interval around the estimate.

To test whether aMK is sensitive to polymorphic weakly adaptive alleles, we used the simulated frequency spectra to estimate the rate of adaptation using published aMK software (*38*). When adaptation is due entirely to strongly adaptive alleles, the estimated value of 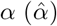 was close to the true value but slightly conservative (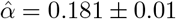; Fig. 1A). As we increased the contribution of weakly-beneficial alleles to *α*, estimates of *α* became increasingly conservative (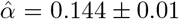 when *α_W_* = 0.1, and 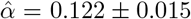 when *α_W_* = 0.2; Fig. 1B-C). Removing polymorphism above frequency 0.5 has been suggested as approach to account for potential biases induced by high-frequency derived alleles, which could be mispolarized in real datasets (*25*). Restricting to alleles below frequency 0.5 produced similar (but conservative) estimates for all three models (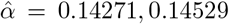, and 0.14264 for *α_W_* = 0.0,0.1 and 0.2, respectively), likely because the frequency spectrum is not strongly dependent on the rate of weakly-beneficial mutation for low-frequency alleles. Lastly, we performed a much larger parameter sweep across *α* values and selection coefficients. We find that *α* estimates become increasingly conservative as the proportion of weakly deleterious alleles increases, and as the strength of selection at beneficial alleles decreases (Fig. S12A & Supplemental Methods). Asymptotic-MK estimates of *α* are only weakly dependent on the distribution of deleterious selection coefficients (Fig. S12).

To better understand why parameter estimates decreased as the proportion of weakly adaptive alleles increased, we performed analytical calculations of *α*(*x*) using diffusion theory (*39,40*). Since we use large sample sizes in our analysis herein, we replace the terms *p_N_*(*x*) and *p_S_*(*x*) in *α*(*x*) with 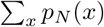 and 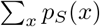 in our calculations, which trivially asymptotes to the same value as the original formulation but is not strongly affected by sample size (see Supplemental Methods). We find that the downward bias in estimates of *α* is due to segregating weakly adaptive alleles, and removing these alleles from the simulated and calculated *α*(*x*) curves would restore the convergence of *α*(*x*) to the true *α* at high frequency (Fig. 1A-C, red curves). In real data, it is not possible to perfectly partition positively selected and deleterious polymorphic sites. Hence, in later sections we focus on using the shape of the *α*(*x*) curve to infer the strength and rate of adaptation under models that include linkage and complex demography.

### Background selection reduces true *α* when adaptation is weak

We have shown that weakly-beneficial alleles may impact aMK analyses by contributing to segregating polymorphism. This presents an opportunity to study whether aMK estimates vary as a function of background selection (BGS) strength. BGS, the action of linkage between deleterious alleles and neutral alleles, reduces genetic diversity in the human genome (*34*) and affects neutral divergence rates (*41*), and is predicted to decrease the fixation probability of weakly adaptive alleles (*33*). Hence, we hypothesized that if adaptation is partially driven by weakly-beneficial alleles in some species, BGS could play a role in modulating adaptation rate across the genome.

To better understand how BGS might affect aMK inference in the presence of weakly-beneficial alleles, we performed analytical calculations and simulations of *α*(*x*) with various levels of BGS. We set *α* = 0.2 in the absence of BGS, and then performed simulations while fixing the rate of adaptive mutations and changing the amount of BGS (ranging from 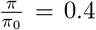 to 1.0, where *π* is neutral nucleotide diversity as compared to the neutral diversity in the absence of linked selection, *π*_0_). We find that when adaptation is strong, BGS has a modest effect on *α*(*x*) and the true value of *α* (Fig. 2A&C), mostly driven by an increase in the rate of fixation of deleterious alleles (Fig. S2E). When adaptation is weak, BGS removes a substantial portion of weakly adaptive alleles and precludes them from fixing, resulting in much stronger dependence of *α*(*x*) on BGS and a substantial reduction in the true value of *α* (Fig. 2B&D and Fig. S2C). Similar to the previous section, estimates of *α* were conservative across all models, but the underestimation was much more pronounced for weak adaptation (Fig. 2C&D).

**Figure 2:**
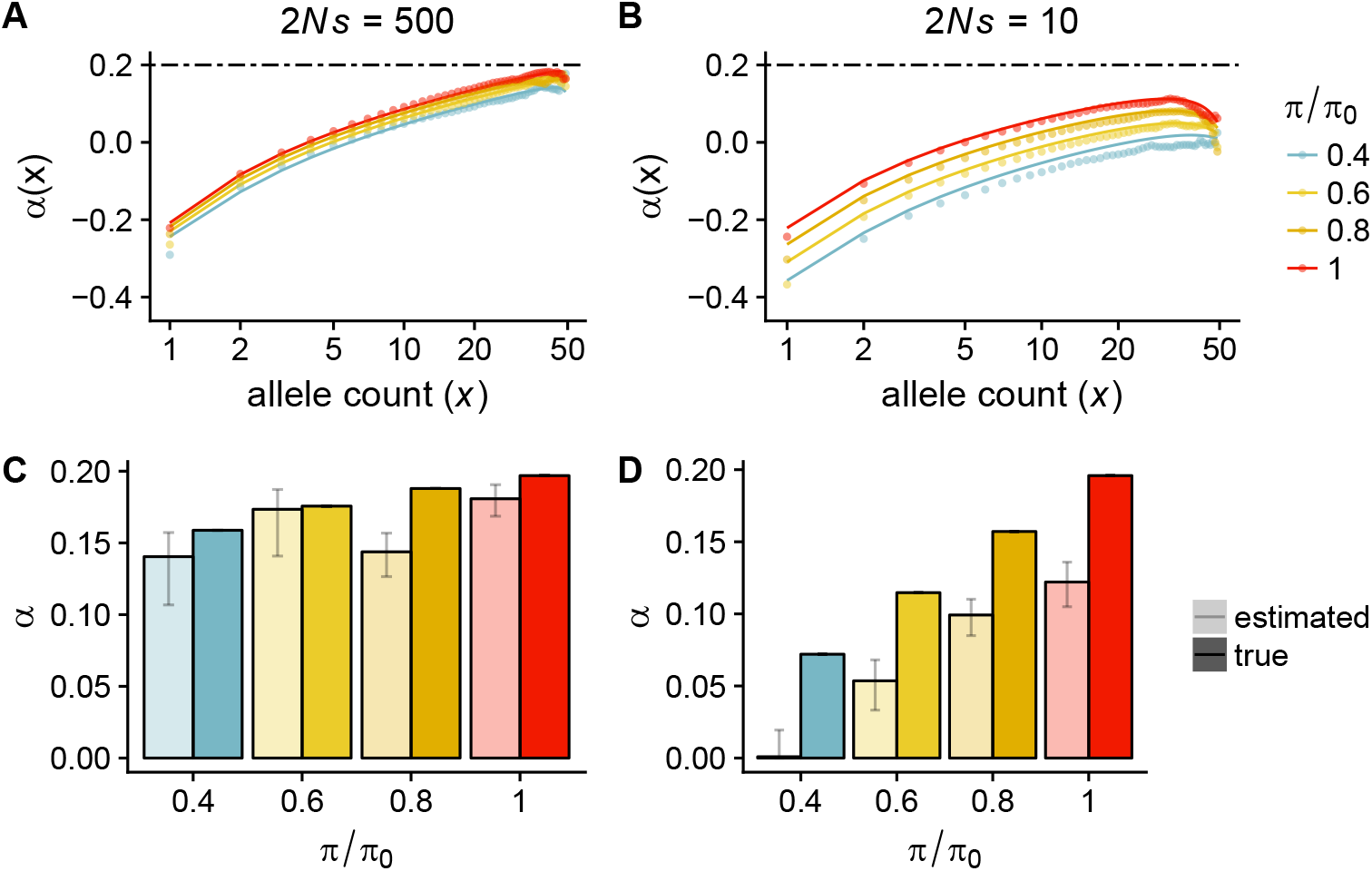
A-B: *α*(*x*) is plotted for various background selection (*π*/*π*_0_) values. In A, adaptive alleles are strongly-beneficial (2*Ns* = 500), while in B they are weakly-beneficial (2*Ns* = 10). The lines represent analytical approximations, while the points represent the results of stochastic simulations. The dashed lines at *α* = 0.2 represent the true rate of adaptation in the absence of BGS. C-D: True (dark colors) and estimated (light colors) *α* for each of the corresponding models in A-B. Panel C corresponds to strong adaptation (2*Ns* = 500) while D corresponds to weak adaptation (2*Ns* = 10). Estimates of *α* were made using existing asymptotic-MK software (*38*). For each parameter combination, we used 2 × 10^5^ independent simulations of 10^3^ coding base pairs each.

### Human adaptation rate is shaped by linked selection

Our modeling results show that *α* is likely to be underestimated when weakly-beneficial alleles contribute substantially to the frequency spectrum, and that background selection may reduce adaptation rate when fitness benefits of adaptive alleles are small. Since BGS is thought to drive broad-scale patterns of diversity across the human genome (*34*), we hypothesized that directly accounting for the action of BGS on adaptation rate could provide new insights into the evolutionary mechanisms driving adaptation. Moreover, the fact that weak adaptation is strongly affected by BGS while strong adaptation is not suggests that strong and weak adaptation could be differentiated in genomic data by comparing regions of differing BGS strengths (from 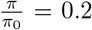 to 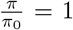). We therefore designed a method to infer *α* while accounting for both BGS and weakly-beneficial alleles.

We developed an approximate Bayesian computation (ABC) approach for estimating *α_W_* and *α_S_* in the presence of BGS and complex human demography (*42*). Briefly, we sample parameters from prior distributions corresponding to the shape and scale of deleterious selection coefficients (assumed to be Gamma-distributed) and the rate of mutation of weakly and strongly-beneficial mutations. We perform forward simulations (*35,36*)of simultaneous negative and positive selection at a coding locus under a demographic model inferred from NHLBI Exome project African American samples (*43*) with varying levels of background selection from *π*/*π*_0_ = 0.2 to *π*/*π*_0_ = 1.0 and the sampled parameter values. We then calculate *α*(*x*) using this simulated data, sampling alleles from the simulations such that the distribution of BGS values in the simulation matches the distribution in the empirical data as calculated by a previous study (*34*). We use *α*(*x*) values at a subset of frequencies x as summary statistics in ABC (specifically, at derived allele counts 1, 2, 5, 10, 20, 50, 100, 200, 500, and 1000 in a sample of 1322 chromosomes). To improve efficiency, we employ a resampling-based approach that allows us to query many parameter values using the same set of forward simulations (see Supplemental Methods). We tested our approach by estimating parameter values (population scaled mutation rates *θ_S_*, *θ_W_*, and the parameters of a Gamma distribution controlling negative selection strength) and quantities of interest (*α_W_*, *α_S_*, *α*) from simulated data. We find that the method produces high-accuracy estimates for most inferred parameters and *α* values (including *α_W_*, *α_S_*, and total *α* – Fig. S6). Some parameter values (particularly those corresponding the the distribution of fitness effects (DFE) over deleterious alleles and mutation rates of beneficial alleles) were somewhat noisily inferred. We find that *α* estimates were not very sensitive to various types of model misspecification (See Supplemental Methods – Robustness analyses), but *α_W_* and *α_S_* are modestly affected by misspecification of the demographic model or the DFE of alleles driving BGS. We term our approach ABC-MK.

We applied ABC-MK to empirical *α*(*x*) data computed from human genomes obtained from the Thousand Genomes Project (TGP) for all 661 samples with African ancestry (*44*). We find strong posterior support for a substantial component of *α* driven by weakly-beneficial alleles (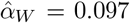; Fig. 3A & see Tab. 1 for area of 95% HPD), as well as posterior support for a smaller component of *α* from strongly-beneficial alleles 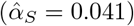. We estimate that the total 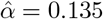, nearly twice the estimate obtained with the same dataset using the original aMK approach (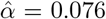, see Supplemental Methods; we note that while our estimate is similar to previous estimates (*23,37*), we use a much larger set of genes in our inference and hence the estimates are not directly comparable). In addition to rates of positive selection, our approach provides estimates of negative selection strength. We find support for mean strength of negative selection of 2*Ns* ≈ −220 (Fig. S9C), which is consistent with recent studies using large sample sizes (*45*) and weaker than earlier estimates using small samples (*37,46*).

**Figure 3:**
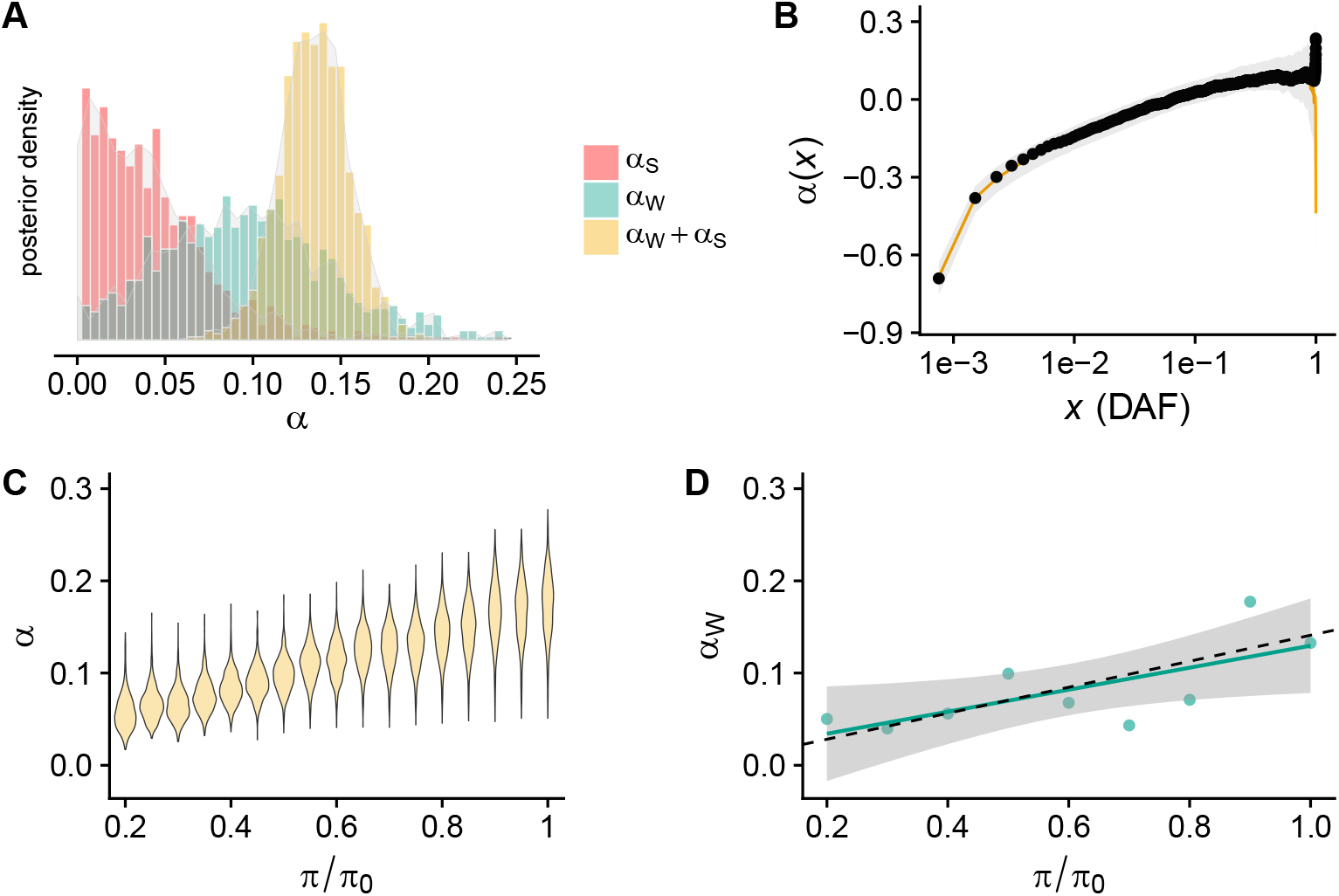
A: Posterior distribution of *α_W_*, *α_S_*, and *α* = *α_S_* + *α_W_* as inferred by applying our ABC approach to 661 samples of African ancestry from the TGP phase 3. B: *α*(*x*) for genomic data (black points) plotted along with the mean posterior estimate from our model (orange line) and 99% confidence interval (gray envelope), as obtained by an independent set of simulations using the posterior parameter estimates. C: Inferred posterior distribution of *α* as a function of BGS strength in the human genome. D: Mean posterior estimates of *α_W_*, as determined by separately fitting the model to alleles from each independent background selection strength bin. A linear model fit to the data (green line) supported statistically significant covariation between *π*/*π*_0_ and *α_W_* (*p*-value=0.0343). The black dashed line shows the predicted change in *α_W_* as a function of *B* given the mean estimate of *α_W_*.

**Table 1:** Table of datasets and inferred values for total adaptation rate (*α*), weak adaptation (*α_W_*) and strong adaptation (*α_S_*). Estimated *α* values represent the mean of the posterior distribution. NS represents the number of nonsynonymous fixations and SYN represents the number of synonymous fixations. Values in parentheses represent the area of 95% highest posterior density.

| Datasets & inferred adaptation rates |  |  |  |  |  |
| --- | --- | --- | --- | --- | --- |
| **Dataset** | **NS** | **SYN** | $\hat{\alpha}$ | $\hat{\alpha}_W$ | $\hat{\alpha}_S$ |
| Whole-exome | 29925 | 38135 | 0.135 (0.096,0.17) | 0.097 (0.0,0.21) | 0.041 (0.0,0.13) |
| VIPs | 6249 | 10309 | 0.224 (0.17,0.28) | 0.098 (0.0,0.24) | 0.126 (0.018,0.26) |
| Non-VIPs | 23676 | 27826 | 0.12 (0.09,0.15) | 0.077 (0.01,0.13) | 0.042 (0.0, 0.09) |

In addition to estimating evolutionary parameters, we sought to better understand how BGS may impact adaptation rate across the genome. We resampled parameter values from our posterior estimates of each parameter, and ran a new set of forward simulations using these parameter values. We then calculated *α* as a function of BGS in our simulations. We find that *α* co-varies strongly with BGS, with *α* in the lowest BGS bins being 33% of *α* in the highest bins (Fig. 3C). Integrating across the whole genome, our results suggest that human adaptation rate in coding regions is reduced by approximately 25% by BGS (Fig. S9D). To confirm that these model projections are supported by the underlying data, we split the genome into BGS bins and separately estimated adaptation rate in each bin. Although these estimates are substantially noisier than our inference on the full dataset, we find that weak adaptation rate decreases as a function of BGS strength in accordance with the model predictions (Fig. 3D). In contrast, estimates of the mean strength of negative selection against nonsynoymous mutations did not covary with BGS strength (Fig. S20). Lastly, to validate that our model recapitulates *α*(*x*) values that we observe in real data, we also used our independent forward simulations to recompute *α*(*x*). We find that our model is in tight agreement with the observed data across the majority of the frequency spectrum. The model and data deviate at high frequency, but both are within the sampling uncertainty (Fig. 3B, gray envelope).

Previous research has shown that virus-interacting proteins (VIPs) have undergone faster rates of adaptation than the genome background (*25*). However, the strength of selection acting on these genes is unknown, and given our BGS results it is plausible that the higher rate of adaptation in VIPs is driven by lower overall background selection at VIPs rather than increased selection pressure for adaptation. In contrast, if pathogens have imposed large fitness costs on humans it is possible that VIPs would support both higher and stronger adaptation rates. We ran our method while restricting to an expanded set of 4,066 VIPs for which we had divergence and polymorphism data available. We found evidence for strikingly higher adaptation rates in VIPs than the genome background (*α* = 0.224) and a much larger contribution from strongly adaptive alleles (*α_S_* = 0.126; Fig. 4). The higher *α* for VIPs cannot be explained by BGS, because VIPs undergo slightly stronger BGS than average genes; the mean BGS strength at VIPs is 0.574, as compared to 0.629 for all genes (in units of *π*/*π*_0_). Taking *α_S_* = 0.126 as a point estimate for the rate of strongly-beneficial substitutions in VIPs and *α_S_* = 0.041 genome-wide, we estimate that 61% of all strongly-beneficial substitutions occurred in VIPs (Tab. 1). Moreover, we estimate that the posterior probability that *α* is greater in VIPs than non-VIPs is 99.97%, while the posterior probability that *α_S_* is greater in VIPs is 88.9% (Fig. 4C). Bootstrap samples of non-VIPs (1,000 replicates) never resulted in *α_S_* estimates as high as those obtained from VIPs (Fig. S19). These results are concordant with the *α*(*x*) summary statistics for VIPs, which had larger values at high frequency alleles than non-VIPs (Fig. 4D). Interestingly, *α*(*x*) is lower for VIPs at low frequency, suggesting increased overall levels of conservation among VIPs (see also Fig. S9, where we find support for stronger negative selection against nonsynonymous mutations in VIPs).

**Figure 4:**
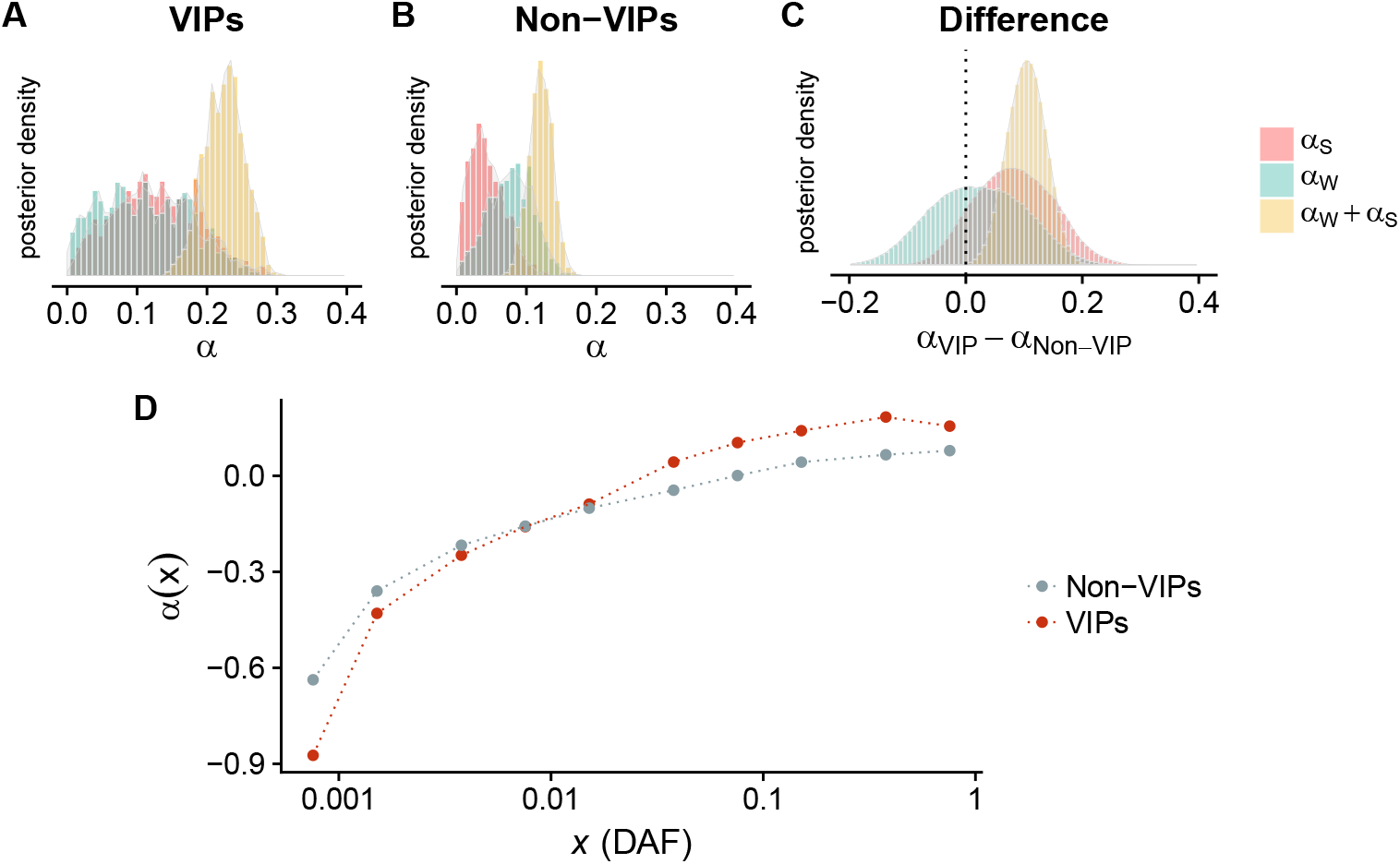
A: Posterior distributions for *α*, *α_W_*, and *α_S_* for virus-interacting proteins (VIPs, 4,066 genes). B: The same quantities for non-VIPs (12,962 genes). C: The posterior distribution of the difference in *α* for VIPs and non-VIPs. D: *α*(*x*) for VIPs and non-VIPs as a function of derived allele frequency *x*, specifically at the values of *x* that we use for statistical inference.

## Discussion

A long-running debate in evolutionary biology has concerned the relative importance of drift and selection in determining the rate of diversification between species (*3, 4, 6, 14*). While previous studies have shown that there is a substantial signal of adaptation in *Drosophila* (*18*), estimates of adaptation rate in humans are much lower (*14*). Here, we extended the classic MK framework to account for weakly-beneficial alleles, and we provided evidence for a large rate of weakly adaptive mutation in humans. We showed that a state-of-the-art approach to adaptation rate estimation that does not account for beneficial polymorphism provides conservative estimates of *α* (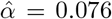 for this data) (*23*), while our method nearly doubles the estimated human adaptation rate (to 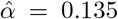). Most of the adaptation signal that we detect is due to weakly-beneficial alleles. Interestingly, virus-interacting proteins supported a much higher rate of adaptation than the genome background 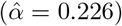, especially for strongly-beneficial substitutions (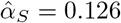 as compared to 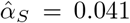 genome-wide). Our results provide an evolutionary mechanism that partially explains the apparently low observed rate of human adaptation in previous studies, and extends the support for viruses as a major driver of adaptation in humans (*25*).

It has long been known that recombination could in principle affect the evolutionary trajectories of both beneficial and deleterious alleles (*33, 47, 48*), and studies in *Drosophila* (*49, 50*) and dogs (*51*) have provided evidence for the effect of recombination on divergence and load. Despite the expectation that recombination could have a strong effect on adaptation in humans, studies have differed on how recombination affects human divergence and polymorphism. One human genomic study explored the ratio 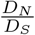 as a function of recombination rate, and found no evidence for an effect of recombination on divergence rate (*11*). Our results may partially explain why 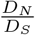 does not fully capture the effect of recombination on divergence in humans. As BGS increases in strength, the rate of accumulation of deleterious alleles increases, while the rate of fixation of weakly adaptive alleles decreases. The two effects partially offset each other, which should reduce the sensitivity of 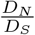 as a tool to detect the effect of recombination on divergence. A more recent study provided evidence that recombination affects the accumulation of deleterious polymorphic alleles (*52*), but did not provide detailed information about the effect of recombination on adaptation. Our results are consistent with the idea that weakly deleterious alleles are predicted to segregate at higher frequencies in regions under strong BGS, and we additionally show that BGS affects the accumulation of weakly-beneficial alleles in humans.

While classic MK approaches estimate only the rate of adaptation, our method extends the MK-framework to provide information about both the rate and strength of selection. Previous approaches to estimating the strength of adaptation have focused on the dip in diversity near sweeping alleles (*31, 49, 53–55*) or have directly inferred the DFE from the frequency spectrum (*20*) – our approach capitalizes on an orthogonal signal of the reduction in fixation rate of weakly-beneficial alleles induced by selection at linked sites. We developed an ABC method to capture this signal, but less computationally intensive methods could also be used – for example, the original aMK approach could be applied in bins of BGS strength. If a substantial proportion of adaptation is due to weakly-beneficial alleles, such an analysis should result in a strong correlation between BGS strength and (potentially conservative) *α* estimates. However, it should be noted that cryptic covariation between gene functions (such as VIPs) and BGS strength could confound such inferences.

We supposed that the main effects of linked selection in humans were due to background selection, but in principle genetic draft could drive similar patterns. Draft is expected to substantially reduce genetic diversity when sweeps occur frequently, and can impede the fixation of linked beneficial alleles (*56, 57*). Previous work has also shown that strong draft can alter the fixation rate and frequency spectra of neutral and deleterious alleles (*23*). We performed simulations of strong draft in 1MB flanking sequences surrounding a gene evolving under natural selection and tested the magnitude of the deviation from theoretical predictions under a model of background selection alone. Consistent with previous work, we observe that draft increases the fixation rate of deleterious alleles and thereby decreases *α* (*23*). However, the effect on *α*(*x*) is only modest at the frequencies that we use in our inference procedure (*i.e*., below 75%), even when the strength and rate of positive selection are much larger than we and others have inferred in humans (although there is a modest deviation around 75% frequency, the highest frequency we use in our inference; Fig. S4C&D). This implies that draft due to selected sites outside genes would have to be much stronger than draft due to positive selection inside exons in order to drive the effects that we infer in the human genome. We note that it is likely that species undergoing both strong, frequent sweeps and BGS (*e.g*., *Drosophila* – see (*31*)), draft will contribute to the removal of weakly-beneficial polymorphism.

Selection has left many imprints on the human genome, with studies reporting signatures of selective sweeps (*55*), soft sweeps (*29*), background selection (*34*), negative selection (*37,46*), and polygenic adaptation (*28*). Still, considerable uncertainty remains about the relative importance of these evolutionary mechanisms, especially as concerns the rate and strength of positive selection. Recent work has suggested that the contrasting adaptation rate estimates of previous studies (*54, 55*) can be reconciled by arguing that most adaptation signals in humans are consistent with adaptation from standing variation (*29*). Our results show that the frequency spectra and patterns of divergence are also consistent with the idea that many adaptive alleles segregate much longer than is expected for a classic sweep, and hence also help to reconcile the results of previous studies.

In addition to determining the rate, strength, and mechanisms of adaptation, there is an ongoing effort to find the biological processes most important for driving adaptation. Previous work has shown that viruses are a critical driver of adaptation in mammals (*25*), but the strength of the fitness advantages associated with resistance to (or tolerance of) infection remain unclear. Our approach clarifies that adaptation to viruses is also approximately three-fold enriched for virus-interacting genes. In contrast, weak adaptation rate was not substantially different between VIPs and non-VIPs, suggesting that weak adaptation may proceed through mechanisms that are shared across proteins regardless of function (for example, optimization of stability). While we have focused on VIPs here due to the expected fitness burdens associated with infection, in future research our approach could be used to investigate adaptation in any group of genes, or extended to partition genes into strong and weak adaptation classes.

The model that we fit to human data does an excellent job of recapitulating the observed patterns in the Thousand Genomes Project data, but we were concerned that several possible confounding factors could influence our results. We showed that seven confounding factors (ancestral mispolarization (*58*), demographic model misspecification (*59, 60*), BGS model misspecification, covariation of BGS and sequence conservation, GC-biased gene conversion (*61*), selection on synonymous alleles (*62*), and misspecification of strongly- and/or weakly-beneficial selection coefficients) are unlikely to substantially influence the results (see Supplemental Methods), but it should be noted that the adaptive process in our model is exceedingly simple, and it is very likely that the evolutionary processes driving diversification are much more complex. We supposed that adaptation proceeds in two categories, weak and strong selection, each of which is described by a single selection coefficient. In reality, adaptive alleles are likely to have selection coefficients drawn from a broad distribution, and adaptation is likely to proceed by a variety of mechanisms, including sweeps (*55*), polygenic adaptation (*28*), and selection from standing variation (*29*). While our results show that BGS shapes adaptation rate across the genome, our method does not differentiate among adaptation mechanisms. We expect that future research will further clarify the relative importance of various selection mechanisms to shaping genomic patterns of diversity in the genomes of humans and other organisms (*10, 63*).

Our method is flexible, and as with the original aMK approach, we showed that the *α* estimates obtained are only minimally affected by demographic uncertainty. It may therefore be an effective tool for providing more accurate estimates of adaptation rate in non-model species that have not been the subject of detailed genomic studies. Despite recent progress, the evolutionary mechanisms that drive the range of diversities observed across species (which could include linked selection, population size, and/or population demography) remain the subject of debate (*12, 13, 16*). Future work using and extending our method, which provides more accurate estimates of adaptation rate when weakly-beneficial alleles contribute substantially to polymorphism, could help to resolve this open question.

## Acknowledgments

We thank Philipp Messer, Raul Torres, Ying Zhen, Christian Huber, Kirk Lohmueller, Zachary Szpiech, Adam Siepel, Yifei Huang, Hussein Hijazi, and our anonymous reviewers for comments that improved the manuscript. We thank Alan Aw, Noah Rosenberg, and members of the Rosenberg lab for helpful discussions. LHU was partially supported by and IRACDA fellowship through NIGMS grant K12GM088033, as well as NIH R01 HG005855 and NSF DBI-1458059 (each to Noah Rosenberg). We thank the Stanford/SJSU IRACDA program for support.

## Supplementary Methods

### Model

We apply a classic directional selection model in which new alleles have selection coefficients s drawn from some distribution over *s*. New mutations arise at rate *θ* = 4*Nμ*, and mutations that are synonymous are treated as neutral whereas nonsynonymous mutations are beneficial or deleterious.

Our ultimate goal is to construct an estimator that jointly infers the rate of adaptation (captured by *α*, which is defined to be the proportion of substitutions that are adaptive) and the strength of selection (*i.e*., the distribution of 2*Ns* values over functional sites). It will be instructive to begin by reviewing the results of Messer & Petrov (*23*), who developed a novel estimator for *α*. Subsequently, we extend their results using analytical theory and simulations to capture information about the strength of selection.

Following earlier work (*18, 23*), we let *d_N_* be the substitution *rate* and we replace *N* in the subscript with *N*_+_, *N*_−_, or *N*_0_ to indicate advantageous, deleterious, or neutral non-synonymous substitutions. When *d_N_* alone appears, it denotes the total rate for all nonsynonymous variants (*i.e*. *d_N_* = *d_N_−__* + *d*_*N*_+__ + *d*_*N*_0__). Analogously, *d_S_* is the substitution rate for synonymous variants, which are assumed to be neutral (and hence do not have additional subscripts).

Consider now the proportion of functional sites that are fixed by positive selection, *α*.

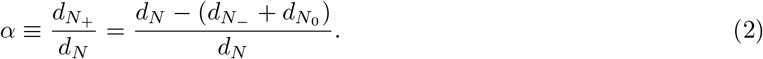

Rearranging, we have

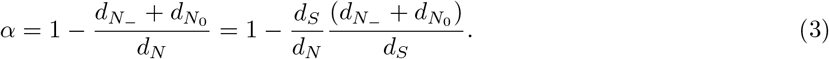

Let the number of observed substitutions be denoted *D*. As noted by (*23*), 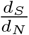 can be estimated from sequence alignments by taking the ratio of *D_S_* and *D_N_*, under the assumption that the observed number of substitutions is proportional to the rate. However, the ratio (*d*_*N*_−__ + *d*_*N*_0__)/*d_S_* is not straightforward to estimate, because the numerator relies on classifying substituted sites by their fitness effects. However, under the assumption that polymorphic sites are rarely selected (because deleterious sites are removed from the population quickly and advantageous sites go to fixation rapidly),

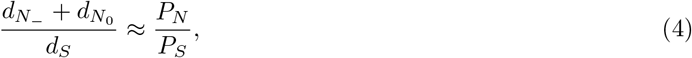

and hence

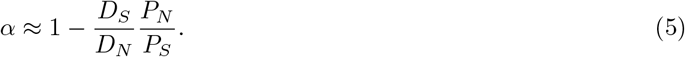

### Assumptions of the MK framework

Approx. 4 implicitly assumes that selected polymorphism is rarely observed. In reality, it is likely that moderately deleterious alleles sometimes contribute substantially to observed polymorphism, especially at low frequency. To guard against this possibility, we can then modify eqn. 5 as

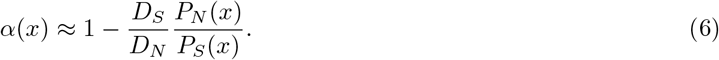

where *P_N_*(*x*) and *P_S_*(*x*) are all non-synonymous polymorphism above frequency *x* and all synonymous polymorphism above frequency *x*, respectively. We note that the original asymptotic-MK approach takes *P_N_* (*x*) and *P_S_*(*x*) as the number of polymorphic sites *at* frequency *x* rather than above *x*, but this approach scales poorly as sample size increases since most common allele frequencies *x* have very few polymorphic sites in large samples. We therefore define *P_N_* (*x*) and *P_S_*(*x*) as stated above since these quantities trivially have the same asymptote but are less affected by changing sample size.

It has been noted that many studies have selected a fixed frequency threshold (say, *x* = 0.15), and removed all polymorphisms below this threshold (*64*). However, if moderately deleterious sites segregate above *x*, then the fixation rate approximation *π*_*N*_−__ ≈ *π*_*N*_0__ is not valid, and *α*(*x*) will be downwardly biased (*64*).

Messer & Petrov (*23*) observed that as the frequency threshold x is increased to be asymptotically close to 1, eqn. 6 asymptotes to the true value of *α*. Intuitively, this is because weakly deleterious alleles (*e.g*., 2*Ns* = −1) can rise to appreciable frequency, but have substantially different fixation probability than neutral sites at all frequencies, meaning that approximation 4 may be poor for all values of derived allele frequency *x* that are substantially less than 1. However, as *x* is increased to be arbitrarily close to the absorbing state at *x* = 1, eqn. 6 approaches the true value of *α* because the probability that a site increases to frequency *x* = 1 − *δ* is a good approximation to the probability that a site fixes for very small values of *δ*.

In most sequencing experiments, there are very few segregating sites with derived allele frequencies close to 1, so simply taking the highest possible value of the threshold frequency *x* results in a very noisy estimator. Hence, Messer & Petrov suggested taking all possible thresholds *x* and fitting an exponential curve to *α*(*x*) (*23*). They showed that when selection is strong, this results in accurate estimates of the adaptation rate *α*.

### Analytical approximation to *α*(*x*)

While the results of Messer & Petrov account for weakly deleterious polymorphic sites, they do not account for the possibility of weakly advantageous sites contributing to *P_N_* (*23*). Here, we use analytical theory to investigate to the quality of the approximation in eqn. 6 when adaptation is weak but occurs at an appreciable rate, such that positively selected mutations occur frequently but fix only rarely. In this section, we assume that the population has constant size, and relax this assumption later with ABC. The calculations in this section proceed similarly to those in previous studies (*22, 65*).

First, we note that while 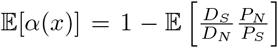 is not straightforward to calculate, the expectation of each quantity on the RHS of eqn. 6 (*i.e*., *P_N_*, *P_S_*, *D_N_*, *D_S_*) is easily calculated from first principles using diffusion theory (*40*). Therefore, we make the first-order approximation

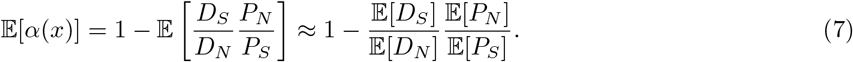

Denoting the distribution of selection coefficients over new mutations as *μ_s_* (multiplied by the underlying mutation rate) and the fixation probability as *π_s_*, the expected number of substitutions along a branch of time *T* in a locus of length *L* is simply

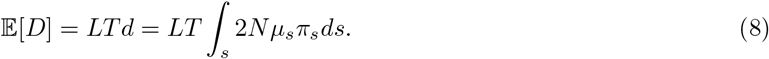

Note that for neutral mutations, where *μ_s_* is non-zero only for *s* = 0 and the fixation probability is given by 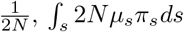 reduces to 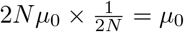.

Likewise, the expected number of polymorphisms above frequency *x* can be calculated from the standard diffusion theory for the site frequency spectrum (*39*), given by

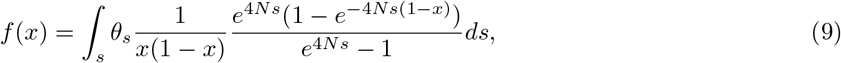

where *θ_s_* = 4*Nμ_s_* is the mutation rate for sites with selection coefficient s. We have assumed that there is no dominance (note that this assumption can be relaxed, but for simplicity we consider only genic selection herein). In a finite sample of 2*n* chromosomes, we must convolute eqn. 9 with the binomial to obtain the downsampled frequency distribution. We deonte the convoluted frequency spectrum as *f_B_*(*x*), defined as the expected proportion of polymorphic sites with allele count equal to *x* in a fixed sample, and note that the total number of polymorphic sites *P*(*x*) in a sample is given by

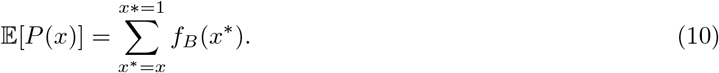

Hence, we can substitute eqns. 8 and 10 into eqn. 6 for *α*(*x*) to make theoretical predictions about the shape of *α*(*x*) as a function of model parameters.

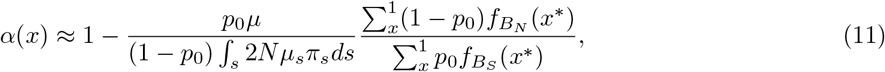

where *f_B_S__* (*x*\*) and *f_B_N__* (*x*\*) are the downsampled site frequency spectra for synonymous and nonsynonymous alleles, respectively, and *p_0_* is the probability that a polymorphic site is synonymous (*i.e*., assumed to be neutral). We developed software that calculates eqn. 11 explicitly for the case of a Gamma distribution of selection coefficients (see next section).

### Gamma distributed selection coefficients

While the previous section did not assume a functional form for the distribution of selection coefficients, in order to perform simulations and inference we supposed that deleterious selection coefficients were Gamma-distributed. Gamma distributions have previously been shown to provide a good fit to human polymorphism data, and have revealed that most nonsynonymous alleles are weakly deleterious, with a long tail of strongly deleterious variation (*37,46*). Additionally, we suppose that advantageous alleles are either strong or weak, such that they are drawn from a point mass distribution with two values (*s_W_* and *s_S_*, where *W* and *S* indicate Weak and Strong).

Replacing *θ_s_* = 4*Nμ_s_* in eqns. 7-8 with a Gamma distribution Γ[*α*, *β*] over selection coefficients (where we have ignored the underlying mutation rate constant, which ultimately cancels out in our calculations), we find that

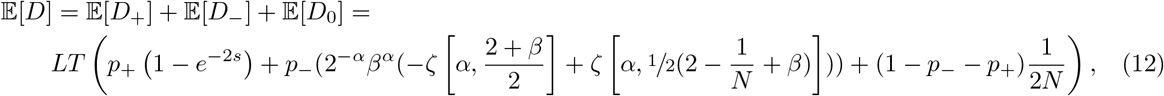

where *p*_+_ is the probability that an allele is deleterious and *p*_−_ is the probability that it is deleterious, and *ζ* is the Riemann Zeta function. The frequency spectra for Gamma distributions of deleterious effects have been previously investigated (*66*).

### Using asymptotic-MK to infer *α*

We used the method of Messer & Petrov (*23*) to infer *α* from the simulated data presented in Fig. 1. This method fits an exponential curve to *α*(*x*) and takes the value of the best-fitting exponential function at *x* = 1 as the inferred value of *α*. In all three panels of Fig. 1, the true rate of adaptation as observed in the simulations is *α* = 0.2, but the component of *α* that consists of weakly adaptive substitutions (*α_W_*) varies from 0 to 0.2 (*i.e*., when *α_W_* = 0.2, all adaptive substitutions are weakly adaptive). To infer *α*, we used published software implementing this method (*38*). The inferred *α* is plotted as a black dotted line in Fig. 1, while the 95% confidence interval is plotted as a gray bar.

We used the default setting for the frequency threshold as provided by the software (*38*), which removes all alleles below minor allele frequency of 10%. When inputting the frequency spectrum for all 661 individuals, we obtained negative estimates of *α*, presumably because there are very few alleles per bin at high frequency in large samples which induces numerical instability. We therefore binned the frequency spectrum into 5% frequency bins in performing the analysis, which resulted in a more stable fit.

In addition to using the previously published software, we also implemented asymptotic-MK in `R` using the function `nls` (nonlinear least squares). We fit a curve of the form *α*(*x*) = *a* + *be^cx^* to alleles between *x* = 0.1 and *x* = 0.9 (*i.e*., the same default range of frequencies used in the previously published software (*38*)). We applied this fitting procedure to predicted *α*(*x*) curves using our analytical approximations. We find that *α* is strongly under-estimated when adaptation is due to weakly-beneficial alleles (Fig. S12A). This result is largely insensitive to the distribution of deleterious alleles – decreasing the mean strength of selection on deleterious alleles did not substantially change the performance of the estimation procedure (Fig. S12B-C). Removing beneficial polymorphism from the frequency spectrum essentially fixes this problem (Fig. S12D-E). Of course, it is not possible to remove the beneficial polymorphisms in real data.

### Background selection & adaptive divergence

Background selection, the action of linked deleterious alleles on patterns of genetic diversity (*67–69*), may also alter the adaptive process. Linked selection reduces the effective population size and hence increases the rate of drift of neutral loci, and may also reduce the efficacy of selection on deleterious alleles and alter fixation rates of both deleterious and positively selected alleles (*33*).

We investigated the impact of background selection on *α* and *α*(*x*) using analytical theory and simulations. We focus on a model in which a coding locus is flanked by loci of length *L* containing deleterious alleles with population-scaled selection coefficient −2*Nt* undergoing persistent deleterious mutation at rate 4*Nμ*_−_. The flanking loci recombine at rate *r* per-base, per-generation. The diversity at the coding locus is decreased relative to its neutral expectation by

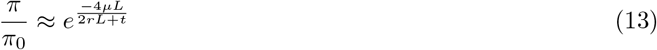

as derived previously (*68, 69*).

The effects of background selection on *d_N_*, *d_S_*, the frequency spectrum, and effective population size have been the subject of much theoretical work (33, 67, 70). It was shown previously (*33*) that the probability of fixation of a positively selected allele under background selection is reduced by a factor *ϕ*, with

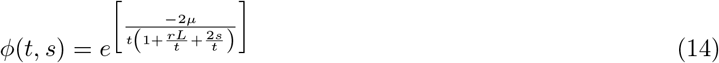

Multiplying across all deleterious linked sites, we find that

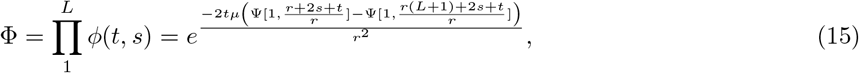

where Φ is the total reduction in fixation probability and Ψ is the polygamma function.

### Testing the analytical theory with simulations

We rigorously tested the theoretical calculations herein using stochastic simulations (*35*). Fig. S1 reports results of background selection simulations, and shows that for a range of expected background selection values calculated with eqn. 13, the expected diversity is in close agreement with values of nucleotide diversity obtained in forward simulations. In our simulations, we solve eqn. 13 for the desired mutation rate in order to obtain the desired reduction in diversity.

We also show that the predicted frequency spectra for positively selected, negatively selected, and neutral alleles are all in close agreement with simulations (Fig. S3), as are the number of diverged sites for neutral (Λ_0_), deleterious (Λ_−_), and beneficial deleterious (Λ_+_) alleles (Fig. S2). Note that the curves in Figs. S2&S3 represent analytical approximations using the results derived herein, and not fits to the data. For these simulations, we assumed that *α* = 0.2, and that the Gamma distribution of deleterious effects is given by a values previously inferred from human nonsynonymous polymorphism with *a* = 0.184 and *b* = 0.000402 (*37*). We relax these assumptions in later sections when performing inference. Python software for performing these calculations and building SFS_CODE command lines is available https://github.com/uricchio/mktest.

### Divergence and polymorphism data

We retrieved the number of polymorphic sites and their allele frequencies in human coding sequences as well as the number of human-specific fixed substitutions in coding sequences since divergence with chimpanzees. Fixed substitutions were identified by parsimony based on alignments of human (hg19 assembly), chimpanzee (panTro4 assembly) and orangutan (ponAbe2 assembly) coding sequences. Human coding sequences from Ensembl v73 (*71*) were blatted (*72*) on the panTro4 and ponAbe2 assemblies and the best corresponding hits were blatted back on the hg19 human assembly to finally identify human-chimp-orangutan best reciprocal orthologous hits. We used the Blatfine option to ensure that even short exons at the edge of coding sequences would be included in the hits. We further used a Blat protein -minIdentity threshold of 60%. The corresponding human, chimp and orangutan coding sequences were then aligned with PRANKs coding sequence evolution model (*73*) after codons containing undefined positions were removed.

For each human coding gene in Ensembl we considered all possible protein-coding isoforms and aligned separately each isoform between human, chimp and orangutan. The numbers of polymorphic or divergent sites are therefore the numbers over all possible isoforms of a human gene (however the same polymorphic or divergent site present in multiple isoforms still counts for one). If a polymorphic or divergent site was synonymous in an isoform but non-synonymous in another isoform, it counted as one non-synonymous polymorphic or divergent site. Only fixed divergent sites were included, meaning that substitutions still polymorphic in humans were not counted as divergent. The derived allele frequency of polymorphic sites is the frequency across all African populations from the Thousand Genomes Project phase 3 (TGP), which comprises 661 individuals spread across seven different subpopulations (*44*). Allele frequencies were extracted from vcf files provided by the TGP for the phase 3 data. In total, 17,740 human-chimp-orangutan orthologs were included in the analysis. Supplemental Data Table S1 provides the number of synonymous and non-synonymous polymorphic or divergent sites for each of these 17,740 orthologs, as well as the allelic frequencies of the polymorphic sites. Polymorphic sites were counted only if they overlapped those parts of human coding sequences that were aligned with chimp and orangutan coding sequences. The ancestral and derived allele frequencies were based on the ancestral alleles inferred by the TGP phase 3 and available in the previously mentioned vcf files (*44*).

Columns in Supplemental Data Table S1 are as follows: First column – Ensembl coding gene ID. Second column – number of non-synonymous polymorphic sites. Third column – respective derived allele frequencies of these sites separated by commas. Fourth column – number of synonymous polymorphic sites. Fifth column – respective frequencies derived allele frequencies of these sites. Sixth column – number of fixed non-synonymous substitutions on the human branch. Seventh column – number of fixed synonymous substitutions on the human branch.

The supplemental table, along with the data that we used to parameterize our model, is available online at https://github.com/uricchio/mktest.

### Background selection data & identifying VIPs

We obtained estimates of background selection strength across the human genome from previous work (*34*) at http://www.phrap.org/othersoftware.html. Since our genetic data was reported in hg19 coordinates, we then used the liftover utility in the UCSC Genome Browser to convert the background selection coordinates from hg18 to hg19 (https://genome.ucsc.edu/cgi-bin/hgLiftOver). We were able to map 17,028 of the 17,740 orthologs to background selection scores. This final set of 17,028 was used throughout the analyses reported in the paper. We classified virus-interacting proteins by using a previously determined set of 4,066 VIPs (*74*).

### Estimating *α* with ABC

#### Motivation for performing ABC

Although we could use analytical theory developed herein to estimate *α*, it is well known that demography also impacts the frequency spectrum of selected alleles (*75, 76*). Some of the impact of recent demography may be attenuated by using the ratio of nonsynonymous to synonymous alleles for inference (since both categories of sites will be affected (*23*)), but failure to incorporate both selection and demography in general can distort inference of both selection and demography (*75*). Since it is not straightforward to calculate the frequency spectrum under generalized models of selection, demography, and linkage (*77–79*), we instead use Approximate Bayesian Computation (ABC) (*42*) to infer selection parameters while accounting for recent demography.

#### Generic ABC algorithm

ABC proceeds by first sampling parameter values from prior distributions, next simulating model outcomes using these parameter values and calculating informative summary statistics, and lastly comparing the simulated summary statistics to observed data. The parameter values that produce summary statistics that best match the observed data form an approximate posterior distribution. An additional linear model can be imposed to correct for the non-0 distance between the simulated and observed summary statistics (*42, 80*).

Here, we follow this generic approach exactly. The main sources of innovation in our method are 1) selecting summary statistics that are informative for estimating *α* values, 2) simulating summary statistics across a range of BGS strengths corresponding to the inferred distribution of BGS strengths in the human genomic dataset, and 3) employing a resampling-based strategy for generating summary statistics that avoids simulating the full model for different parameter combinations.

#### Overview of our ABC approach

Our ABC approach proceeds in three main steps. First, we run forward simulations with a fixed DFE over nonsynonymous alleles (*37*), a known demography inferred from human genomic samples, and a fixed DFE over deleterious alleles flanking the central coding locus. Second, we used biased resampling of alleles from these simulations to calculate summary statistics that correspond to a wide range of nonsynonymous allele DFEs with varying rates of adaptation and exactly the same number of sampled variants and distribution of *B* scores as the observed data (where *B* is the strength of BGS from a previous study (*34*)). Lastly, we supply 10^6^ sets of summary statistics sampled from our prior distributions into a published ABC software framework (*80*) to infer parameters from real datasets. Below we describe each of these steps in more detail.

We simulate a sample of 661 individuals (the same number of samples as the African continental group in the TGP phase 3 data) under a demographic model incorporating an expansion in the African ancestral population and recent exponential growth (*43*). Within each coding region, we suppose that the distribution of deleterious effects is given by a Gamma-distribution with *a*_0_ = 0.184, *b*_0_ = 0.000402, which were previously inferred as the strength of negative selection in another study using human coding sequences (*37*) (note that the mean strength of negative selection is given by 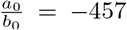, but the distribution is very heavy-tailed with a substantial contribution from weakly deleterious variants). SFS_CODE simulates sequences under the explicit 64-codon genetic code - using this model, approximately 75% of new mutations in coding regions are nonsynonymous. We additionally simulate positive selection with *θ_W_* = 7.8 × 10^−6^ for weak adaptation and *θ_S_* = 2.6 × 10^−7^ for strong adaptation (see below for rationale on selecting these values).

We repeated these simulations over a range of background selection strengths, from 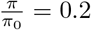 to 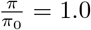 in increments of 0.05. Each simulation replicate consists of a central coding locus of 10^3^ bp flanked on each side by 2 × 10^5^ bp of non-coding loci. We use eqn. 13 to compute the mutation rate for deleterious alleles in the flanking sequences such that the desired reduction in diversity is obtained. We simulate 10^5^ genes of 10^3^ bp in length, for 10^8^ total bp of sequence for each BGS value.

We seek to infer four parameters, which we draw from prior distributions – in particular, *θ_W_* = 4*Nμ_W_*, the mutation rate for weakly-beneficial alleles, *θ_S_* = 4*Nμ_S_*, the mutation rate for strongly-beneficial alleles, and *a* and *b*, the parameters of the Gamma distribution controlling the distribution of deleterious alleles. Since each of these parameters are fixed in our original round of simulations, we resample alleles from the simulated data to reflect the desired combination of selection parameters (see below for resampling details). Using the resampled frequency spectra, *D_N_*, and *D_S_*, we calculate *α*(*y*) for values of *y* in 1, 2,4, 5,10, 20, 50, 200, 500,1000, where *y* is the derived allele count and the frequency x in *α*(*x*) is given by *x* = *y*/2 × 661. Lastly, a linear model is imposed to correct for the non-0 distance between the summary statistic values in the simulations as compared to the observed data. We use previously published software to perform this inference step (*80*).

We additionally infer the *α* values (*α*, *α_W_*, and *α_S_*) – while these are not parameters of the model, they can be inferred in the same ABC framework since they can easily be calculated for any given parameter combination. As priors, we suppose that *θ_W_* is uniform on [0, 7.8 × 10^−6^] and *θ_S_* is uniform on [0, 2.6 × 10^−7^]. We chose these values because at the top of the range, *α_W_* = 0.4 and *α_S_* = 0.4 when the distribution of deleterious effects is given by a Gamma distribution with *a*_0_ = 0.184, *b*_0_ = 0.000402, which were previously inferred in another study using human coding sequences (*37*). We supposed that *a* and *b* might deviate from their previously inferred values by up to a factor of 4 above or below their previous estimates, and hence we sampled exponents *a*_fac_ and *b*_fac_ uniformly on [-2,2] and we let *a* = *a*_0_2^*a*_fac_^ and *b* = *b*_0_2^*b*_fac_^. Hence our prior for a and b are centered at *a*_0_ and *b*_0_, but can vary to allow substantial flexibility in the distribution of deleterious effects. In all of our simulations, we suppose that strongly advantageous alleles have 2*Ns* = 500 and weakly advantageous alleles have 2*Ns* = 10, and we rescale the simulated ancestral population size to *N* = 500. We use a large s approximation for calculating the fixation probability of strongly advantageous alleles by treating the adaptive allele trajectory as a Galton-Watson process (*57*).

Code that we used to simulate these models is available at https://github.com/uricchio/mktest. Note that we designed this software to run on the Stanford HPC cluster, Sherlock – adapting it to run in other computing environments would require further modifications. We suggest that parties interested in using the software contact the authors for assistance in installing it and applying it.

#### Resampled summary statistics & validation

We resampled polymorphic sites from our set of forward simulations with *a* = *a*_0_, *b* = *b*_0_, *θ_W_* = 7.8 × 10^−6^, and *θ_S_* = 2.6 × 10^−7^ to compute summary statistics for ABC. The underlying idea of these resampling simulations is that given a fixed strength of BGS, the allele frequency spectrum can be approximated by selecting alleles in proportion to their mutation rate given the model parameters relative to the parameter values that were used in the original set of simulations. For example, if we suppose that alleles with *s* = 0.001 have a mutation rate of *θ* = 10^−5^ in the original forward simulations but *θ* = 10^−6^ in the resampling simulations, then we resample such alleles at a rate that is 10% of their representation in the original simulations.

For polymorphic positively selected sites, we resample with replacement from the simulated frequency spectra by selecting adaptive polymorphic sites with probability proportional to 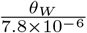 and 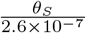 for weakly and strongly-beneficial alleles, respectively. We resample negatively selected alleles with replacement from the frequency spectrum, but we adjust the sampling probability in proportion to the probability that a polymorphic site with selection coefficient s is observed at frequency *x* given the parameter values a and b using the analytical expressions developed in the previous sections. We also analogously adjust the simulated number of fixation events at nonsynonymous along the simulated branch. We confirmed that our resampling-based approach provides the appropriate frequency spectra by comparing simulated resampled frequency spectra to forward simulations performed in SFS_CODE for a subset of parameter values at the boundary of our prior distributions (Fig. S11).

To capture the impact of background selection, we ran the original forward simulations with varying amounts of BGS in 5% bins ranging from 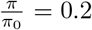 to 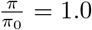 and the same parameter values as above. To calculate summary statistics corresponding to the desired parameter values, for each allele in our TGP dataset we obtained an estimate of BGS strength at the corresponding locus (*34*) and we sampled a polymorphic allele randomly from the frequency spectrum of the simulated BGS bin that is closet to the observed value. We excluded all sites with *B* < 175 (*i.e*., 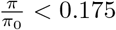) from the inference for computational efficiency, because simulating large reductions in diversity requires high mutation rates of deleterious alleles in the flanking sequences. We pool all of the simulated polymorphic sites to calculate the *α*(*x*) summary statistics corresponding to the model parameters. Open source software implementing our approach is available by request and will be posted online.

We tested our ABC approach by simulating a large dataset of parameter values and matched summary statistics, and then masking a subset of the parameter values. We tested our ability to infer the masked parameter values using the remaining summary statistics for 100,000 replicates. We plot the results of this experiment in Fig. S6, where we summarize the inferred parameter value as the mean of the posterior distribution. We find that the method returns accurate and unbiased estimates for most quantities of interest, although we find that the parameter b controlling the distribution of deleterious effects is somewhat noisily estimated.

### Summary of robustness analyses

Although our model explains the observed *α*(*x*) data very well, we were concerned that several possible confounders might also produce similar patterns. We focused on seven sources of confounding, namely 1) ancestral state uncertainty, 2) covariation of BGS and sequence conservation, 3) demographic model misspecification, 4) misspecification of the strength of selection at sites driving background selection, 5) biased gene conversion, 6) selection acting on synonymous alleles, and 7) misspecification of the strength of selection at adaptive variants.

Ancestral mispolarization could confound our results if some loci with high-frequency derived alleles in our dataset are in fact loci with low frequency derived alleles. Mispolarization can have similar effects on the frequency spectrum as positive selection, and has been identified as a possible source of bias in selection inference (*58*). To limit the effects of ancestral state uncertainty on our analysis, we only use the summary statistics used in our ABC to frequencies at or below 75%, which are much less susceptible to the effects of mispolarization (*58*). Our results are therefore unlikely to be affected by mispolarization.

Covariation between BGS and sequence conservation could also be a potential source of bias in our approach. If negative selection is stronger per site in genes under strong BGS, then the frequency spectrum and rate of fixation of weakly deleterious alleles will also vary as a function of BGS strength (denoted *B* – note that a large *B* corresponds to weak BGS), potentially confounding our results. To test the hypothesis that sequence conservation and *B* covary, we computed the average “rejected substitution” score (RS, as determined by the GERP algorithm (*81*)) on a gene-by-gene basis as a function of *B*. *RS* scores represent the number of substitutions per site that have been rejected due to negative selection, and increase with the strength of negative selection. We found a slight negative correlation between *B* and *RS*, almost entirely driven by genes with *B* > 875 (Fig. S10). While this correlation is consistent with our model (since we expect more substitutions due to weak adaptation in regions with low BGS), it could also be due to the confounding covariation. To eliminate the potential confounding effect of covariation between *B* and sequence conservation, we repeated our ABC-based inference procedure after removing all genes with *B* > 875 from the analysis. If our signal were driven by this covariation rather than a true effect of weakly advantageous alleles, we would expect our parameter estimates to change substantially in this experiment, in particular by increasing the mean strength of selection against deleterious nonsynonymous alleles. In contrast, we observe almost no change in the estimated negative selection parameters (Fig. S9), and when we estimated negative selection strength separately for each BGS bin, we did not observe any covariation with *B* (Fig. S20).

Another possible confounder is demographic model misspecification. Selection and population demography both affect the frequency spectrum, and hence failure to accurately account for both demography and selection in inference procedures can result in biases (*59, 60, 75, 82–84*). Although the aMK framework may avoid some of these issues by directly comparing nonsynonymous and synonymous alleles (*23*), both of which are subject to the same demography, we nonetheless tested for demographic biases. To test the effects of model misspecification, we varied the size of the expansion event in the African ancestral population by sampling parameter values from the 95% confidence interval of a previous demographic model (*85*) that was built using TGP sequences (see Supplemental Methods). We simulated under these models with larger or smaller than expected bottlenecks, and used summary statistics of our “misspecified” model to perform inference of the selection parameters. We find that *α* is still inferred very accurately, although a subset of simulations resulted in over-estimates of *α* when the true expansion was much larger or much smaller than the expected expansion (Figs. S7&S8). We also observed modest biases in *α_W_* and *α_S_*, with *α_W_* under-estimated when the magnitude of the expansion is over-estimated and over-estimated when the expansion is under-estimated (Fig. S7&S8), but the vast majority of inferred total *α* values fell close to the diagonal in both cases. These results suggest that our main results are robust to recent demographic uncertainty, although slight quantitative biases in *α_W_* and *α_S_* could be induced by demographic model misspecification.

Misspecification of the strength of selection acting on alleles driving BGS could also cause bias in our inferences. We supposed that the mean strength of selection against alleles inducing BGS was γ = 2*Ns* = −83, which reflects a mixture of previous estimates of the strength of selection against polymorphism in human coding (*37*) and conserved non-coding (*86*), weighted by the percentage of the genome that is composed of each type of element. If the true strength of selection driving BGS was much smaller or much larger, we might change the expected dependency of *α*(*x*) on *B*. In essence, if γ is closer to 0, BGS should have a smaller effect on the fixation rate of weakly-beneficial alleles. We therefore considered a range of γ values from −10 to −100 – consistent with expectations, we find that weaker selection against BGS alleles induces *α*(*x*) to vary less markedly as a function of BGS strength, but the effect is very modest (Fig. S13).

To further address the possibility that misspecification of the BGS DFE could affect our results, we also repeated our entire inference pipeline with three additional distributions of fitness effects for the BGS alleles (a weak DFE with 2*Ns* = −10, a strong DFE with 2*Ns* = −500, and a gamma-distributed DFE that mixes strong and weak alleles as inferred by Boyko et al (*37*) – see Fig. S14). These varying DFEs had almost no effect on the inferred value of *α*, but note that the gamma DFE resulted in slightly lower estimates of *α_W_* but slightly higher estimates of *α_S_*. Accordingly, we conclude that our results are not strongly dependent on the strength of selection against alleles driving BGS, but misspecification of the BGS DFE could result in slight biases in the weak and strong components of *α* that are similar in magnitude to misspecification of the demographic model (Figs. S7&S8). Lastly, for each of the BGS DFEs that we considered, we checked explicitly that purely non-adaptive simulations could not fit the data. We took the subset of simulated summary statistics that approximately match the empirical *α*(*x*) data at low frequency (*i.e*., fall within 0.1 of the lowest-frequency data point) and additionally have very low adaptation (simulated *α* < 0.01). We plot these summary statistics along with the real data (Fig. S15). The simulated summary statistics fall below the real data at all values of *x* at high frequency.

We also supposed that biased gene conversion (BGC) could be a confounder in our results. BGC can mimic positive selection by favoring the fixation of weak to strong mutations (*87*). We therefore recomputed *α*(*x*) using the 661 TGP samples after removing all the weak to strong mutations and fixations from the dataset. We find that the empirical *α*(*x*) curve is not substantially affected by the removal of weak to strong sites at frequencies that we use for ABC (Fig. S5), suggesting that BGC is unlikely to affect our inferences.

In response to a suggestion by multiple reviewers, we also considered the impact that selection against synonymous alleles might have on our results. Recent studies have suggested that synonymous alleles are likely to be subject to weak negative selection (*62, 88*), which would violate an assumption of the MK test framework. We tested the robustness of our results to this assumption by calculating the analytical expectation of *α*(*x*) curves using a DFE over synonymous alleles that mimics recently inferred distributions from human genomic data. In particular, Huang & Siepel found that 70.5% of mutations were effectively neutral, while 26% were moderately deleterious and 3.5% were strongly deleterious (*62*). We modeled this by mixing neutral alleles (70% of alleles) with a Gamma distribution inferred from conserved non-coding sites (*86*) – under this distribution (gamma parameters *a* = 0.0415 and *b* = 0.00515, which has a mean value of 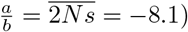) 27% of alleles are weakly or moderately deleterious (|2*Ns*| < 10) and 3% are strongly deleterious (|2*Ns*| > 10). We also considered an *ad hoc* distribution with gamma parameters a = 0.1 and b = 0.1, which has mean 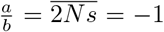 and 29% weakly deleterious alleles and 1% strongly deleterious alleles – we made this choice since weakly deleterious alleles are more likely to be problematic for the MK framework, due to their potential to segregate for long times. Our calculations proceed by simply replacing the neutral terms in eqn. 11 with a mixture of neutral and deleterious distributions.

In Fig. S17, we show *α*(*x*) summary statistics for these synonymous DFEs as compared to purely neutral alleles, which differ only subtly from each other. Crucially, when adaptation is absent, selection on synonymous alleles does not induce false adaptation signals (Fig. S18). The results of these calculations suggest that selection against synonymous alleles would have to differ substantially from current estimates in order to strongly affect our inference. In practice, cryptic selection against synonymous alleles may affect inference through the application of non-equilibrium demographic models, which are often inferred from synonymous alleles or other putatively neutral variants. Multiple studies have now found that demographic inference can be biased by linked selection or direct selection (*23, 59, 60, 83, 89*). More work will be needed to better understand how such inference errors will affect adaptation rate estimation in the MK framework.

Lastly, we considered the impact that misspecification of the strength of positive selection for alleles in the strongly-beneficial and weakly-beneficial categories might have on inference. We performed analytical calculations for a variety of selection coefficients to compare their *α*(*x*) summary statistics. For strong alpha, there is almost no difference in the computed summary statistics in the absence of BGS and a small difference when BGS is strong (Fig. S16). This is the expected behavior, since strongly-beneficial alleles rarely contribute to segregating polymorphism and are only very weakly affected by BGS. For weak alpha, there are small differences in the values of the summary statistics for each of the *α*(*x*) curves when BGS is absent - when BGS is strong most adaptive alleles are removed and the difference between different selection coefficients is diminished. This suggests that while strongly- and weakly-beneficial alleles have qualitatively different behaviors, we will have little power to infer the full DFE over beneficial alleles from these summary statistics. This does not preclude the possibility that more complex DFEs could also fit the summary statistics (see Discussion section in main text).

### Genetic draft

Our modeling uses a diffusion approximation to dynamics of allele frequency shifts that accounts for background selection but not draft. If genetic draft (*i.e*., the impact of linked positive selection on the frequency trajectories of linked alleles) also drives systematic variation in diversity genome-wide, then this approximation may break and invalidate some of the assumptions of our modeling (*23*).

To test the sensitivity of our results to genetic draft, we compared simulations with and without genetic draft to our theory for a range of selection strengths and rates. We simulated a gene under simultaneous negative and positive selection, flanked by 1 MB sequences. We compared models with and without BGS, and with and without draft, for a range of parameter values. We set *α* = 0.4 within the gene, and supposed that 5% of the flanking sequence was a potential target for positive selection that was both as strong and as frequent as that within the gene.

Consistent with earlier results (*23*), we find that draft can decrease *α*, likely by increasing the rate of fixation of weakly deleterious alleles and/or interference between strongly-beneficial alleles (*56, 57*). However, even in the extreme scenario where adaptation is driven by very strongly advantageous alleles with 2*Ns* = 2000 and *α* = 0.4, we observe only a modest departure from the expectation in the absence of draft at the frequencies that we use in inference, all but one of which are below 37% frequency (Fig. S4). This suggests that our inference should be only modestly affected by draft, and only in regions of the genome experiencing strong, recurrent sweeps.

### Simulations for demographic model misspecification

We tested the impact of demographic model misspecification by sampling “worst-case” parameters from the 95% confidence interval of a previous study that fit a maximum-likelihood demographic model to TGP sequences (*85*). The maximum likelihood estimates from this model for the ancestral human population size and expanded population size are *N_A_* = 7, 300 and *N_AF_* = 12, 300, respectively. We supposed that the largest possible expansion would correspond to the 2.5% quantile estimate of *N_A_* and the 97.5% quantile estimate of 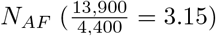, while the smallest possible expansion would correspond to the 97.5% quantile estimate of Na and 2.5% quantile estimate of 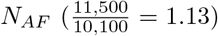. We then ran simulations under our model, sampling parameters from the same prior distributions as described above, and generated summary statistics. We then attempted to infer the parameters that were used to generate the summary statistics using our misspecified demographic model. Results of this experiment are shown in Fig. S7 & Fig. S8, and are described in the main text.

## Supplemental figures

**Figure S1:**
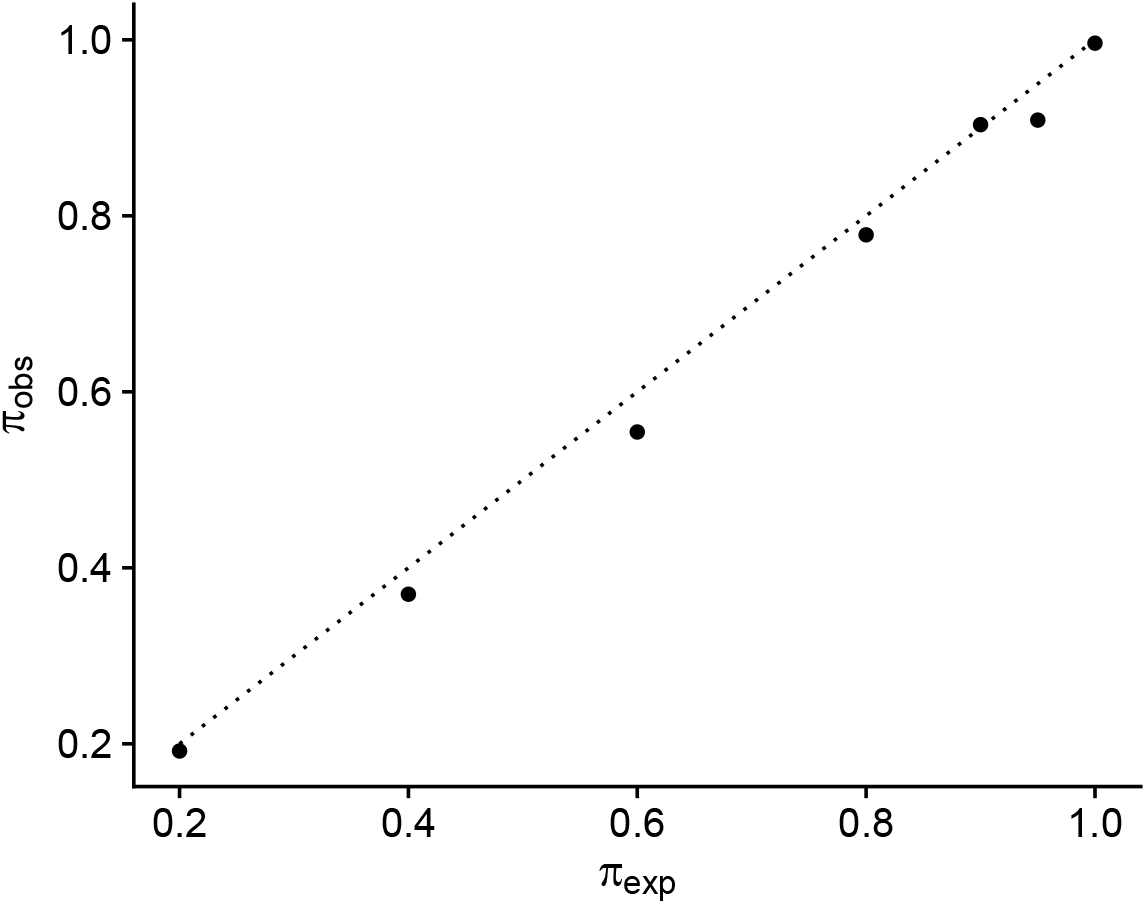
Simulated(*π*_obs_) vs expected (*π*_exp_) nucleotide diversity for simulations performed in SFS_CODE. The expected value was calculated using the model of Hudson & Kaplan (*68*).

**Figure S2:**
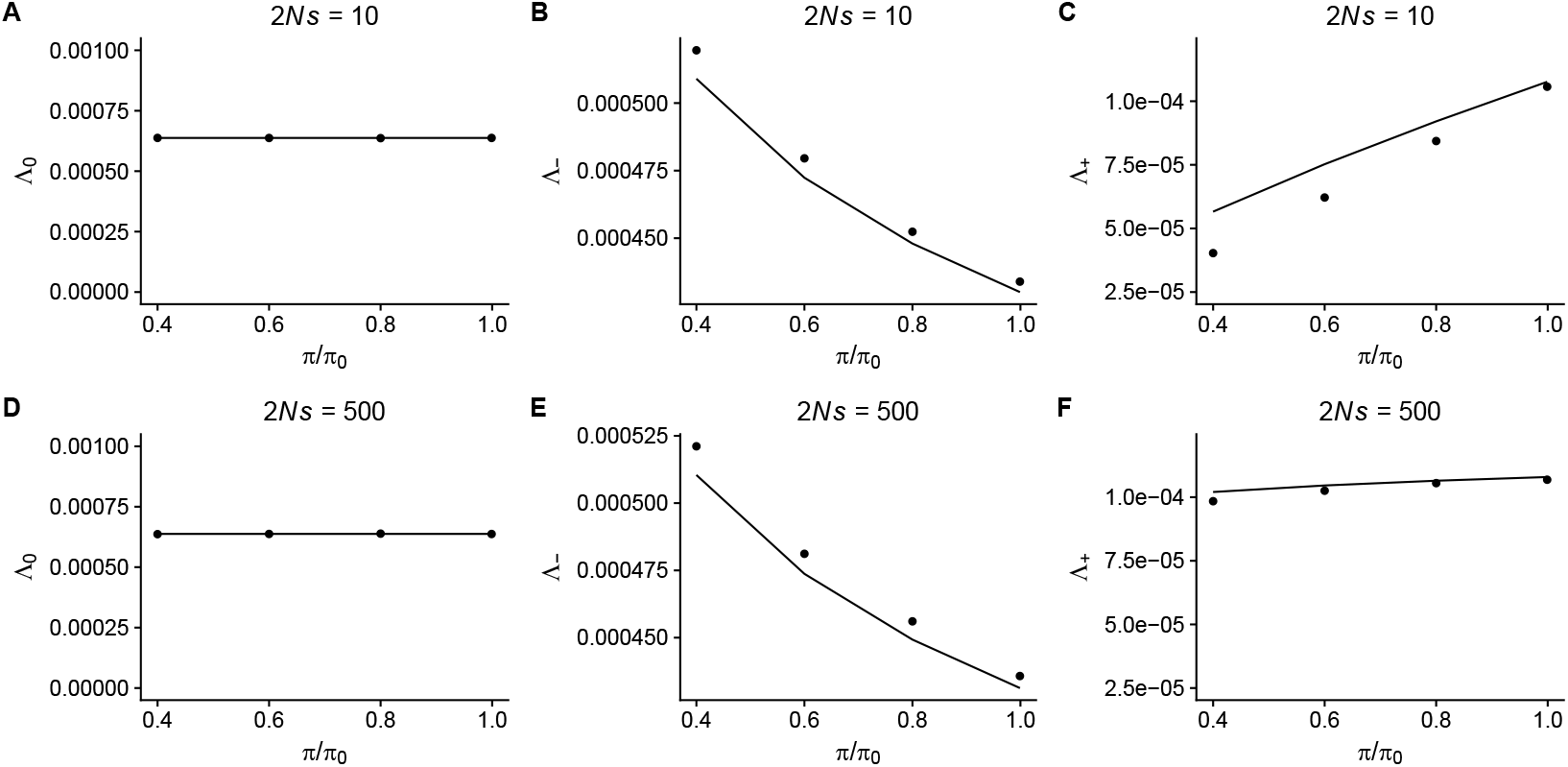
Simulated (points) and expected (lines) fixation rates for neutral, negatively selected, and positively selected alleles. Eqns. for the expected fixation rates are given in the supplemental text. The top row represents results in the context of weakly-beneficial adaptation (2*Ns* = 10), while the bottom row represents strongly-beneficial adaptation (2*Ns* = 500).

**Figure S3:**
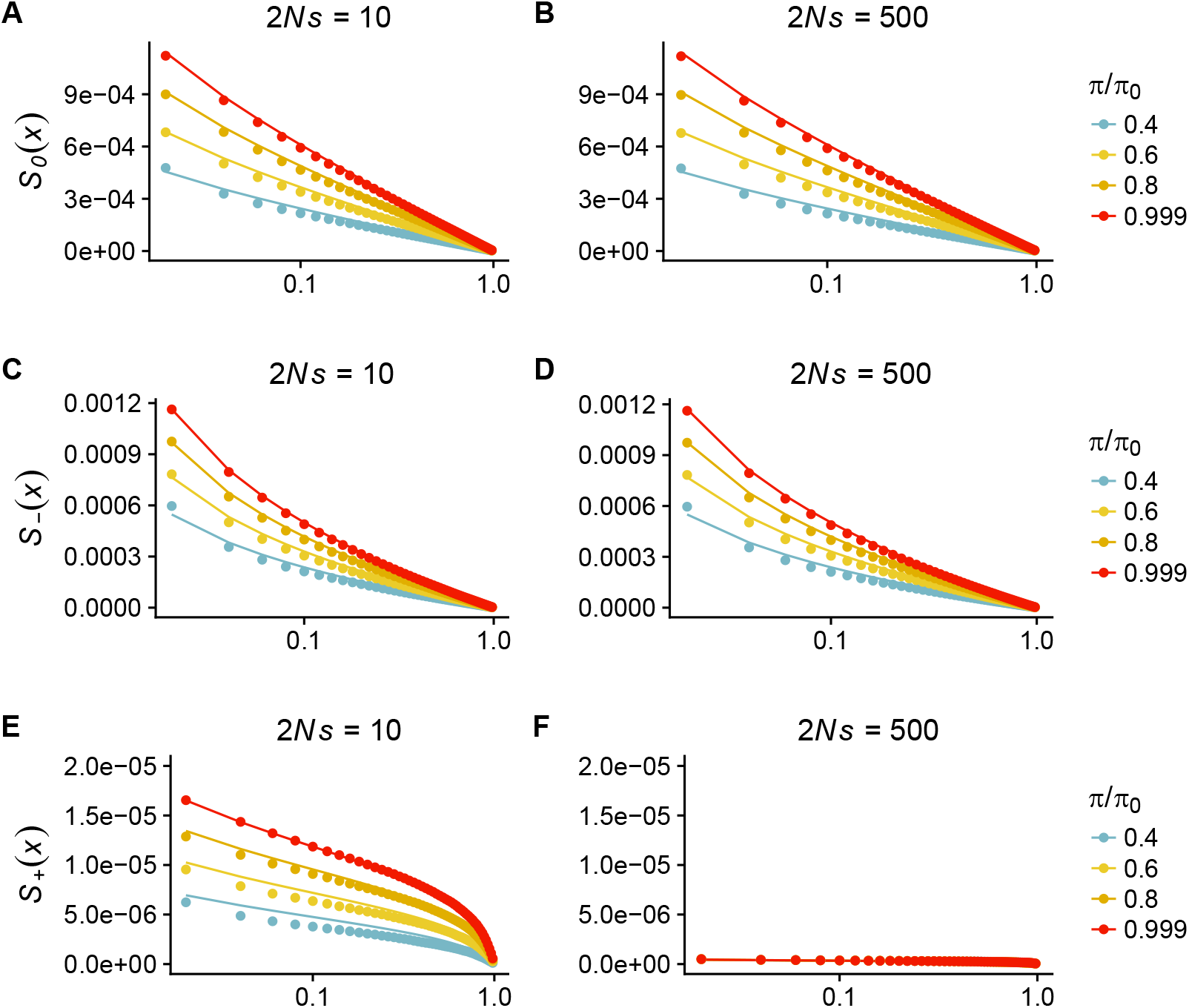
Simulated (points) and expected (lines) frequency spectra for neutral, negatively selected, and positively selected alleles. *S*(*x*) is the number of alleles above frequency *x*.

**Figure S4:**
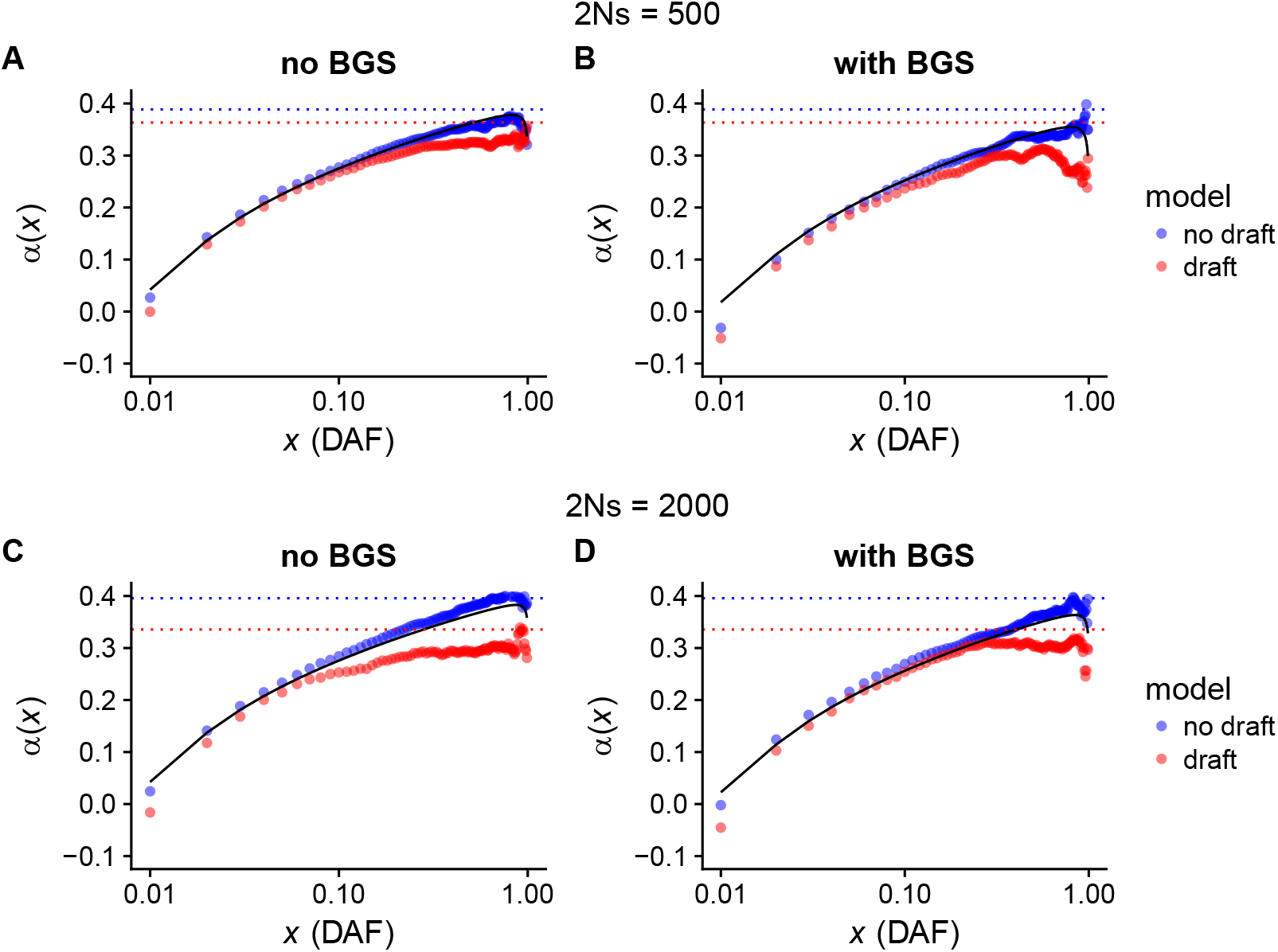
Comparison of simulations with and without genetic draft. In all simulations we set *α* = 0.4, and suppose that 5% of the sequence in the 1MB flanking a gene is subject to recurrent sweeps. The black line shows the theoretical expectation from eqn. 11.

**Figure S5:**
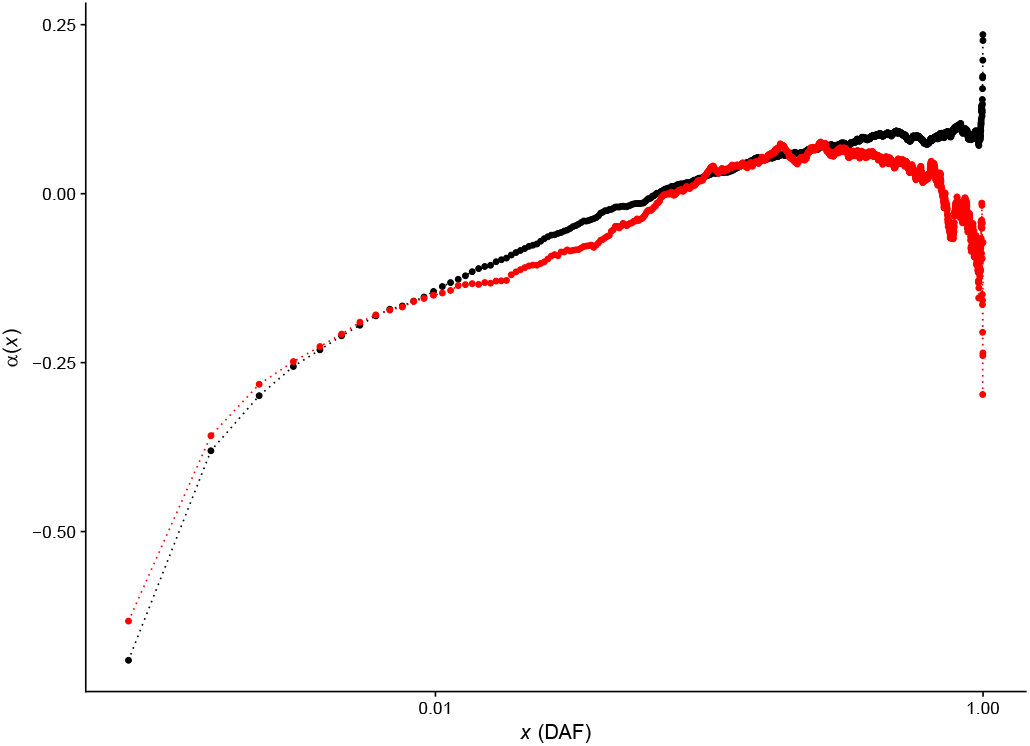
Comparison of *α*(*x*) computed from TGP samples for all sites (black) and with weak to strong sites removed (red).

**Figure S6:**
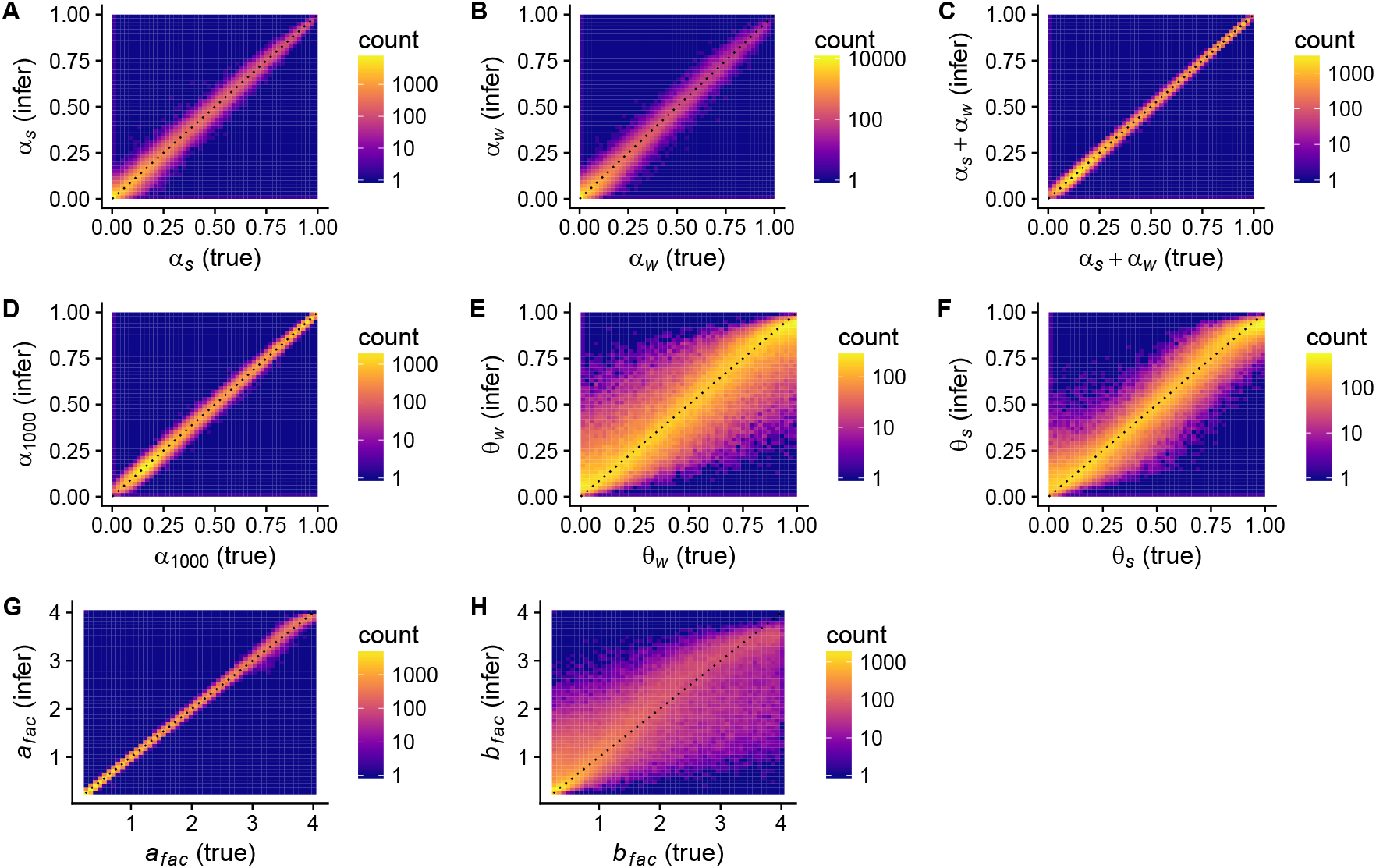
Performance of our parameter estimation for all of parameters and quantities that we infer. In each panel, the true parameter value is plotted on the *x*-axis, while the inferred value is plotted on the *y*. The diagonal is plotted as a dashed black line. The inferred value is summarized as the mean of the posterior distribution. Each plot contains 100,000 simulations.

**Figure S7:**
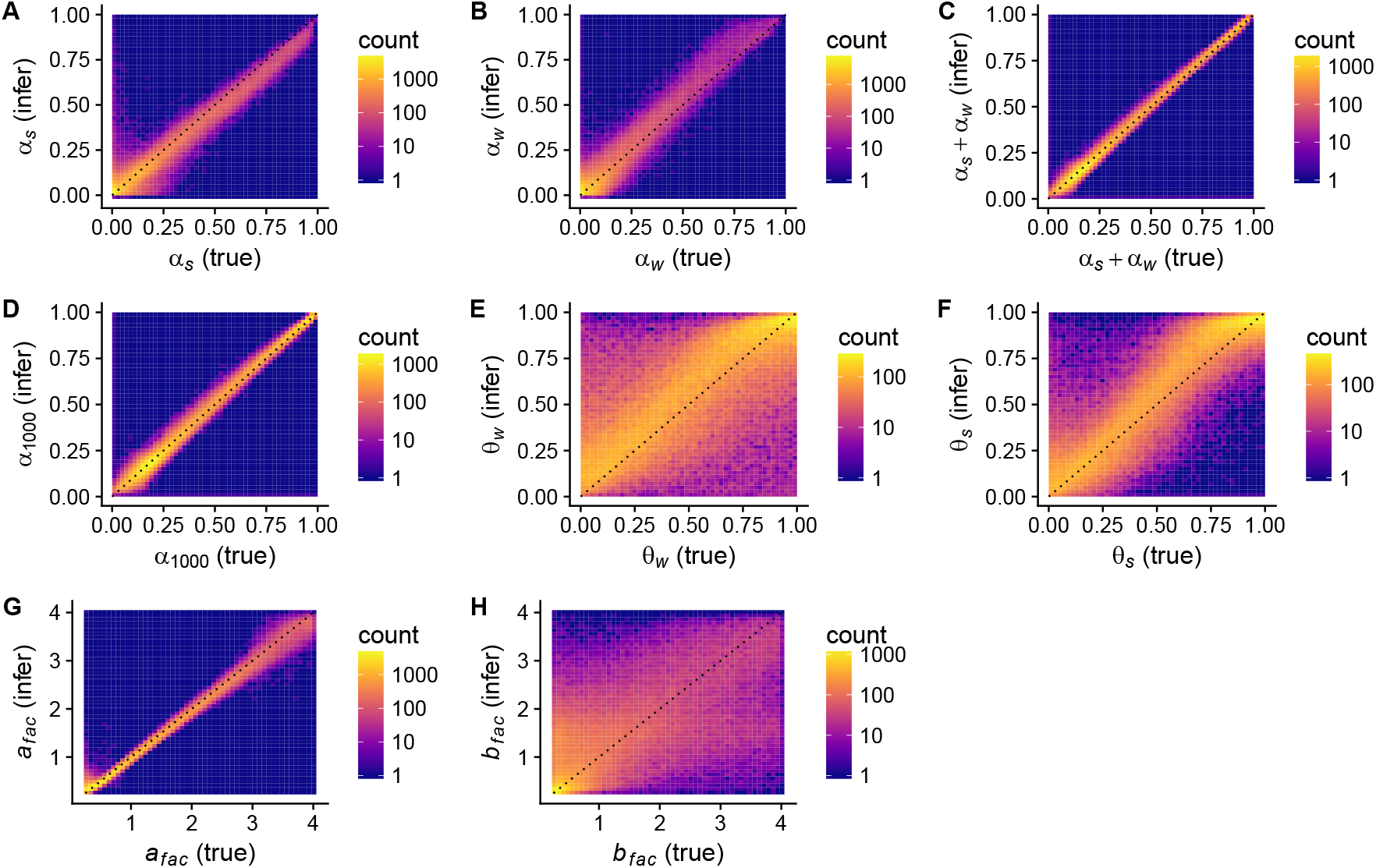
Performance of our parameter estimation for all of parameters and quantities that we infer, in the case when the true model has an ancestral expansion event that is ≈ 2 larger than the model used in the inference procedure. In each panel, the true parameter value is plotted on the *x*-axis, while the inferred value is plotted on the *y*. The diagonal is plotted as a dashed black line. The inferred value is summarized as the mean of the posterior distribution. Each plot contains 100,000 simulations.

**Figure S8:**
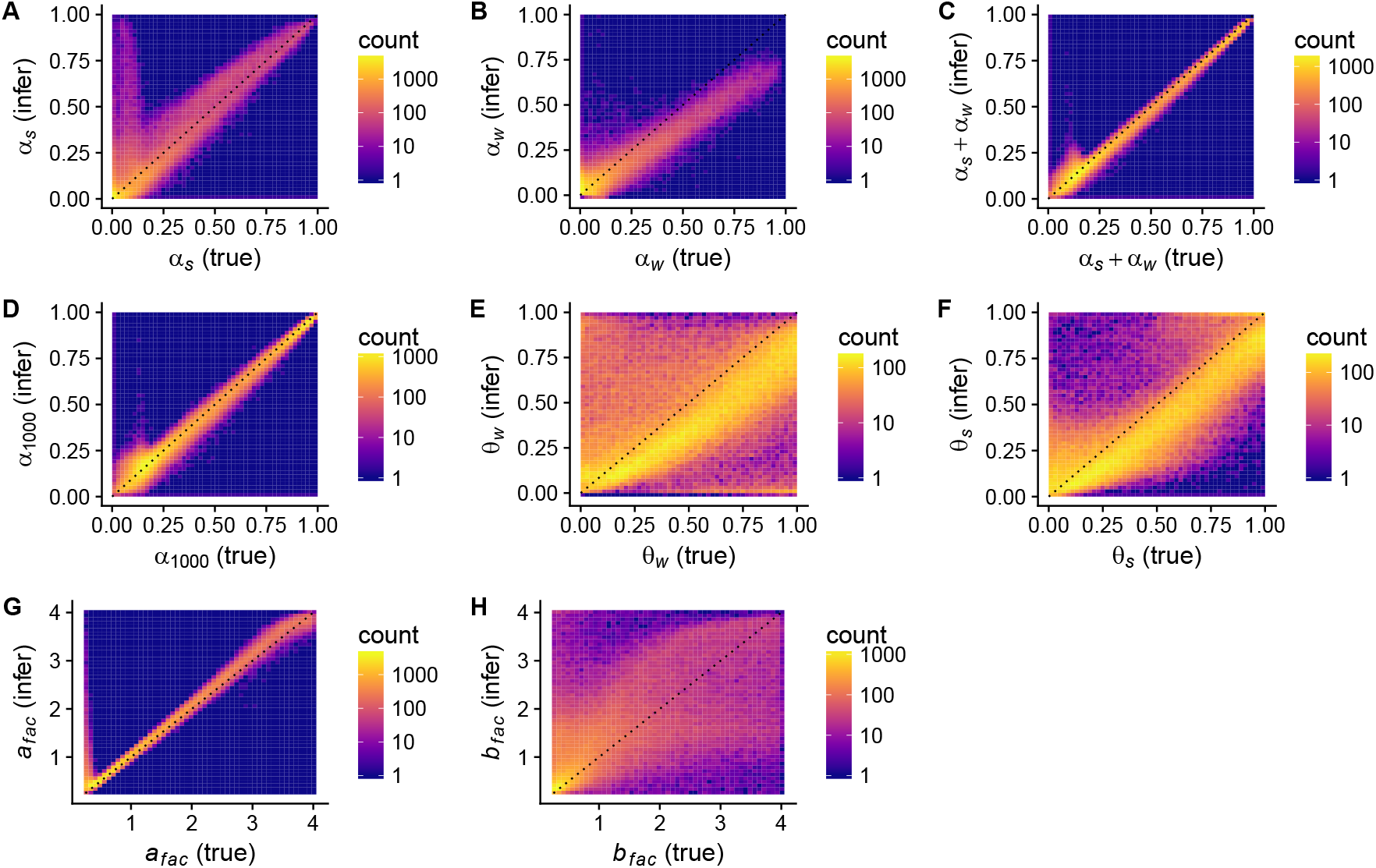
Performance of our parameter estimation for all of parameters and quantities that we infer, in the case when the true model has an ancestral expansion event that is 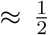 as large as the model used in the inference procedure. In each panel, the true parameter value is plotted on the *x*-axis, while the inferred value is plotted on the *y*. The diagonal is plotted as a dashed black line. The inferred value is summarized as the mean of the posterior distribution. Each plot contains 100,000 simulations.

**Figure S9:**
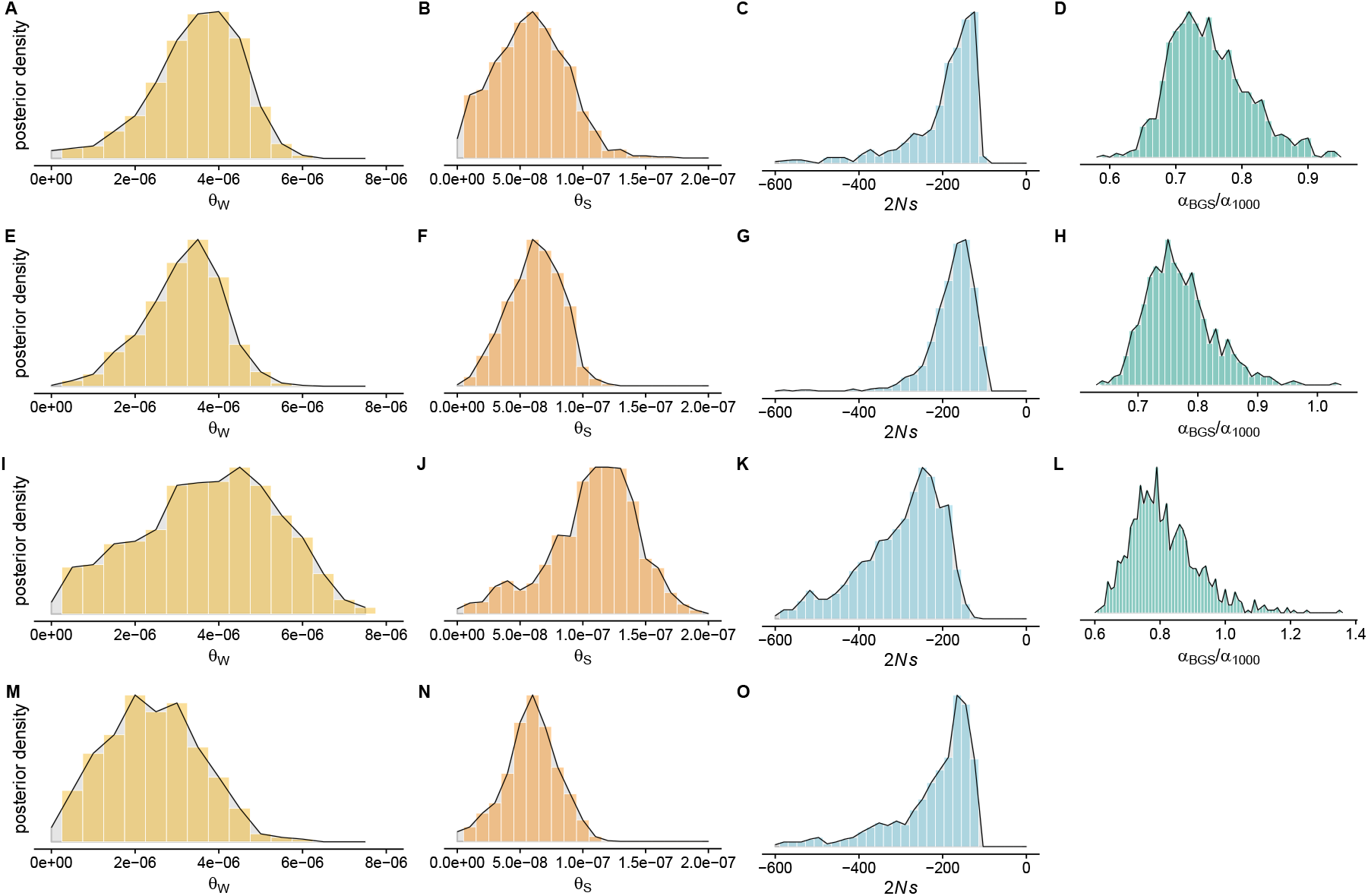
A-D. Posteriors for *θ_W_*, *θ_S_*, the mean strength of negative selection (2*Ns*), and the ratio of *α*_BGS_ (the estimated value of *α* in humans after accounting for BGS) to *α*_1000_ (the value of *α* for regions of the genome with *B* = 1000, *i.e*., regions unaffected by BGS) as predicted by our model. E-H: The same quantities, as inferred using only genes that were not classified as VIPs. I-L: The same quantities, as inferred using only genes that were classified as VIPs. N-O: The same quantities, as inferred using only genes with *B* < 875. We do not infer *α_BGs_*/*α*_1000_ in this row because genes with *B* ≈ 1 are not included in this analysis.

**Figure S10:**
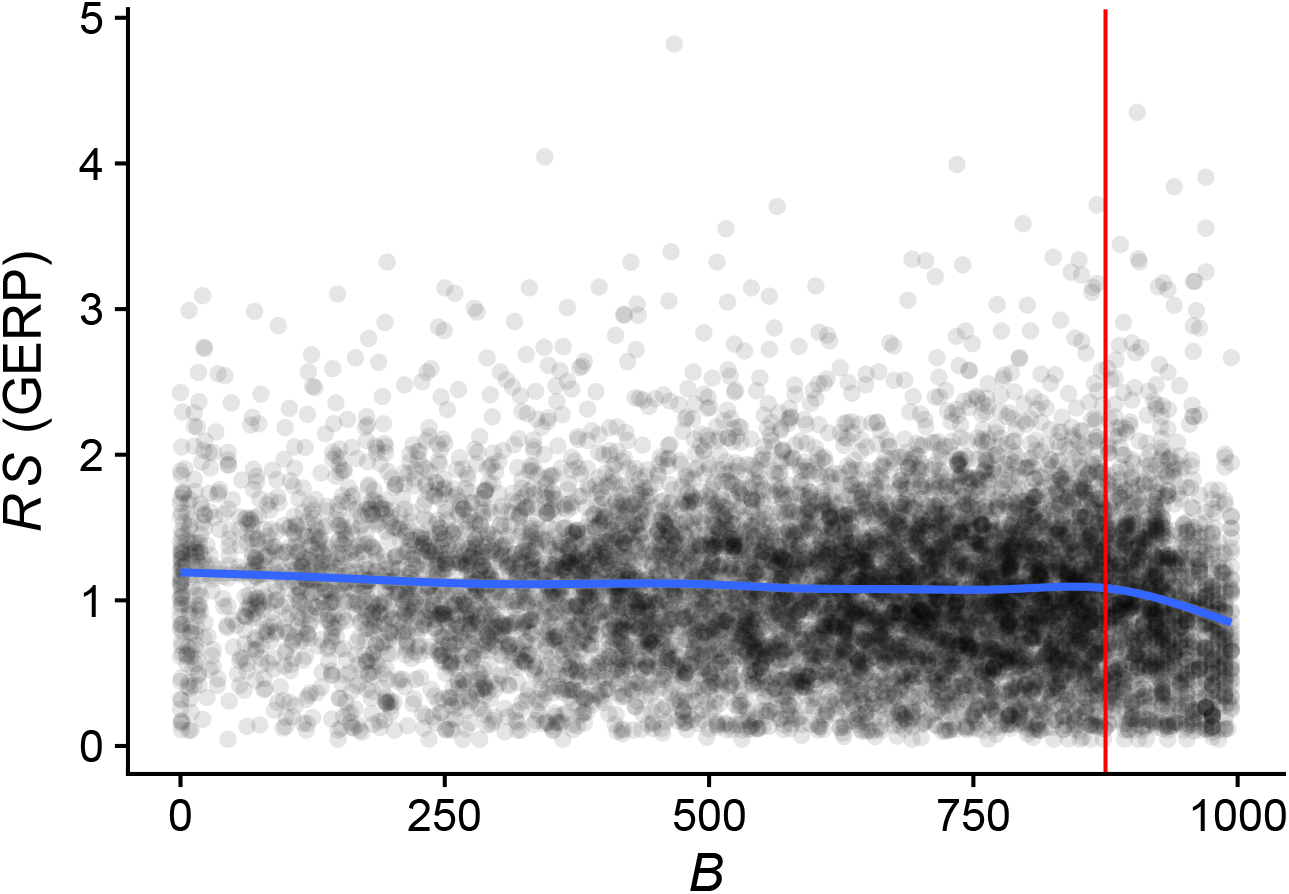
The relationship between BGS (*B*) and average sequence conservation (*RS*) for ≈ 10,000 genes for which we were able to obtain estimates of both quantities. The blue line is fit to the data using `geom_smooth` in `ggplot2`, while the red line is plotted at *B* = 875. Most of the negative correlation between *B* and *RS* is driven by alleles with *B* > 875. Note that *B* is defined in previous work (*34*), and is equivalent to 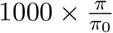.

**Figure S11:**
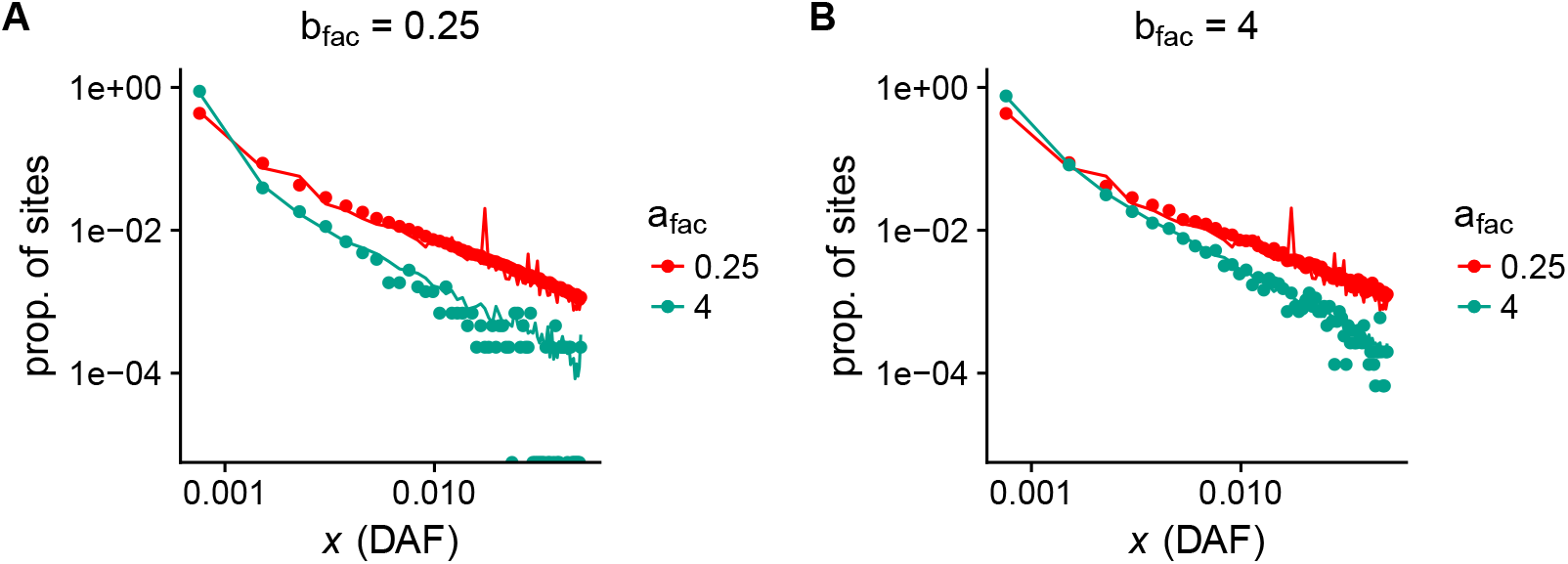
We compare simulated frequency spectra obtained with SFS_CODE (points) to frequency spectra that we obtained using our resampling-based approach (lines) for a range of parameter values corresponding to the strength of negative selection. We observe good agreement between the approaches. One downside of the resampling based approach is that stochastic fluctuations in the dataset from which resampling is performed are replicated across different samples (*e.g*., the spike at ≈ 0.015 is replicated in both A and B).

**Figure S12:**
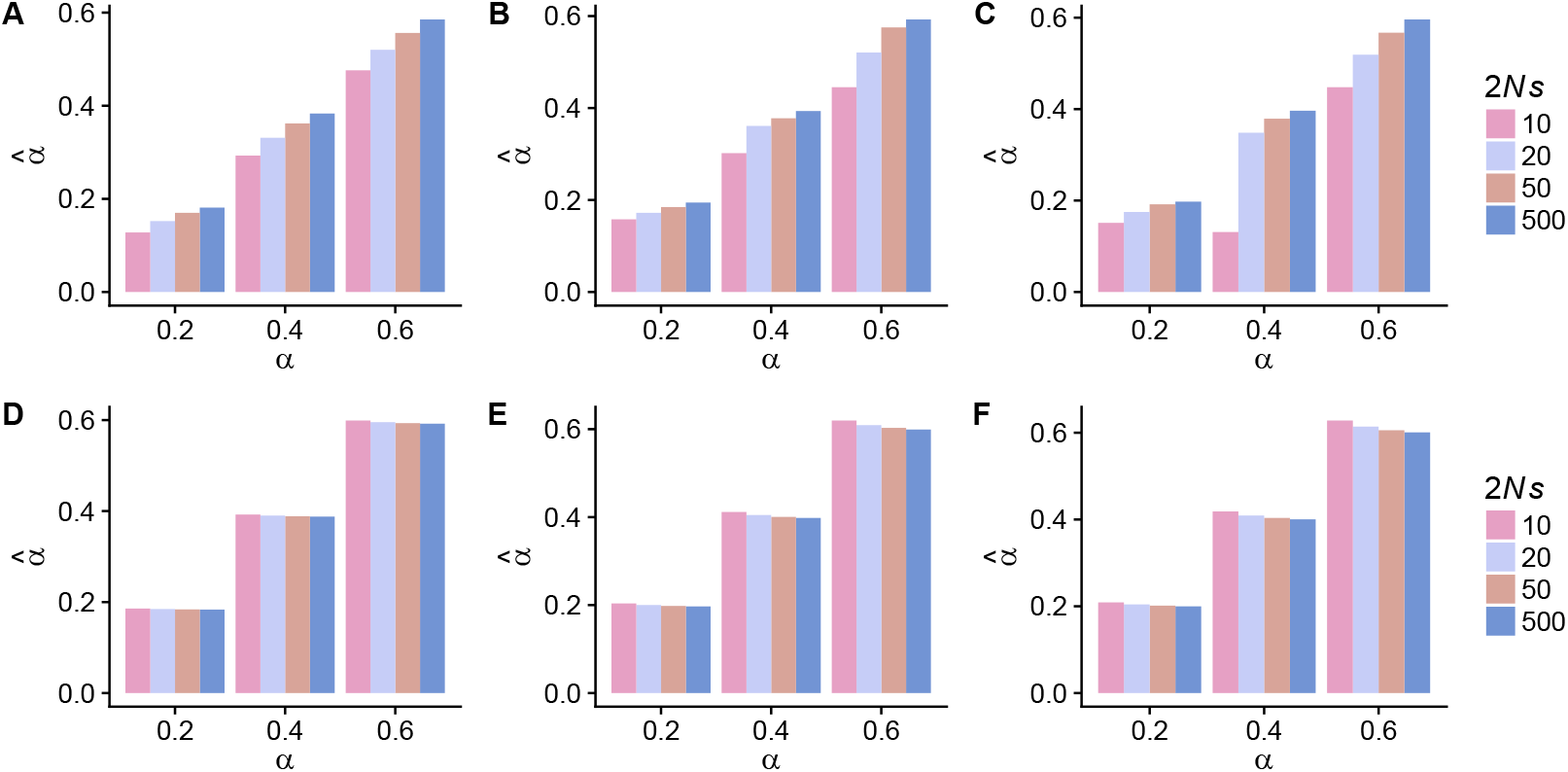
We plot estimates of adaptation rate (*α*) using asymptotic-MK as a function of true *α* for a range of 2*Ns* values of adaptive alleles (colors) and a range of deleterious selection coefficient distributions (each panel is a different distribution of deleterious effects). A&D correspond to the distribution of deleterious effects inferred in (*37*) (which has a mean value of 2*Ns* = −457), while B&E have a mean value of 2*Ns* = −114 and C&F have mean 2*Ns* = −22. In A-C, all alleles are used in the estimation procedure, while in D-F we exclude positively selected alleles from the calculation.

**Figure S13:**
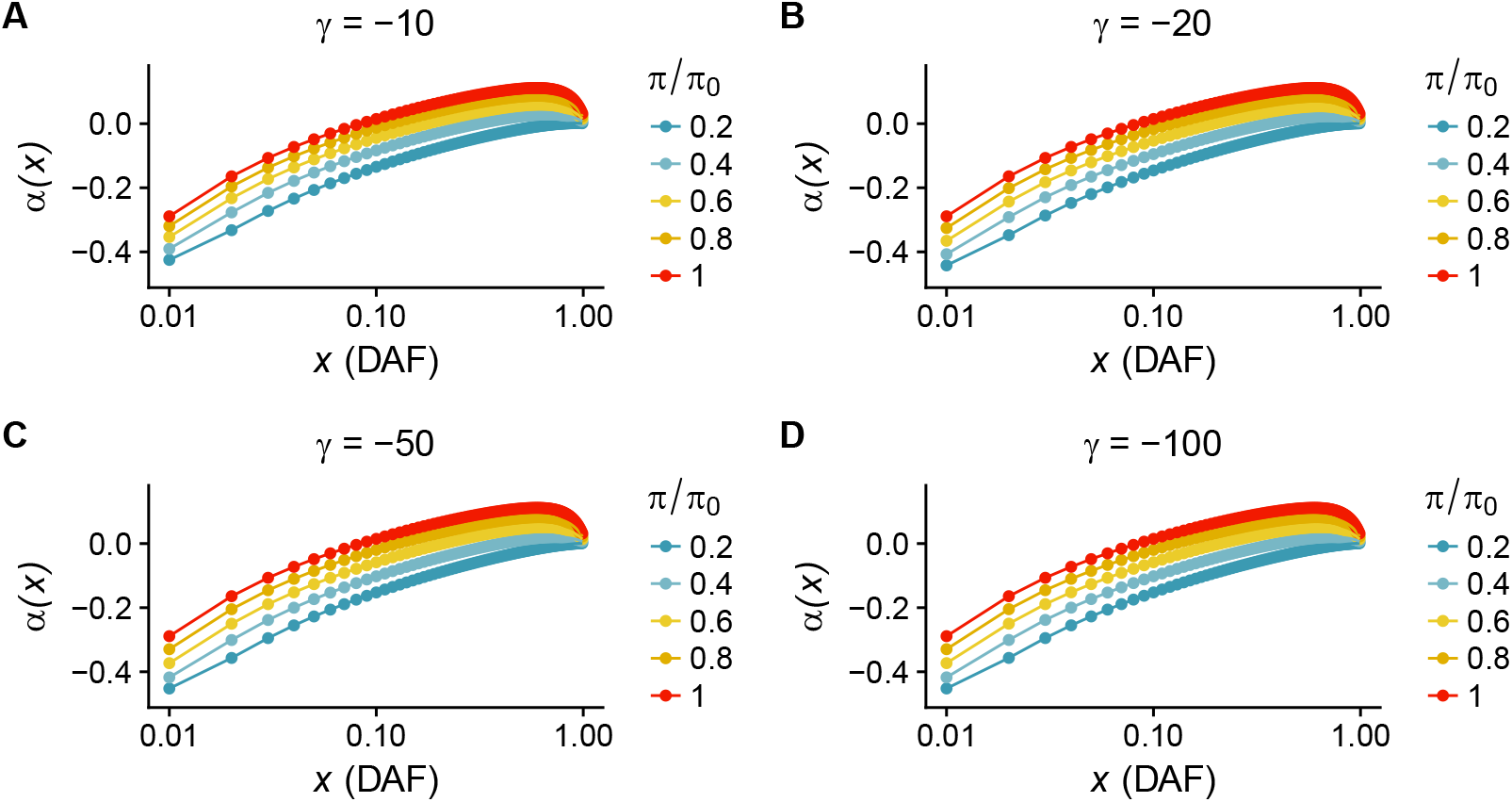
*α*(*x*) as a function of DAF for a range of selection strengths (γ) on alleles driving BGS. Each curve represents a different value of 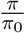. In each panel, the strength of selection on adaptive alleles is 2*Ns* = 10.

**Figure S14:**
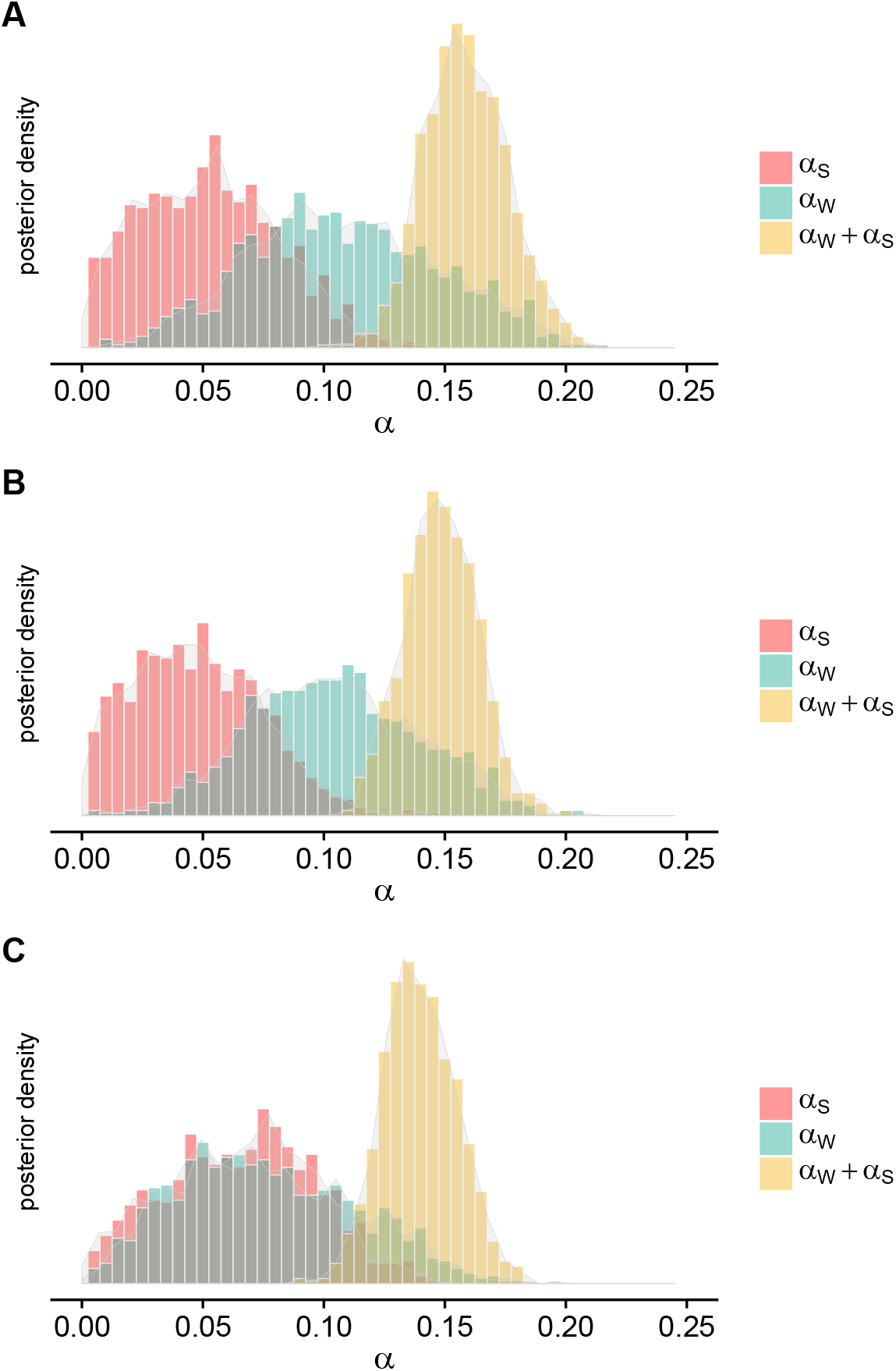
Inferred posterior distributions of *α_S_*, *α_W_*, and *α_S_* + *α_W_* for three different DFEs of alleles driving BGS. In A, alleles in flanking regions around genes have 2*Ns* = −10, in B 2*Ns* =—500, and in C 2*Ns* is a gamma-distributed mixture of weakly deleterious and strongly deleterious alleles (*37*).

**Figure S15:**
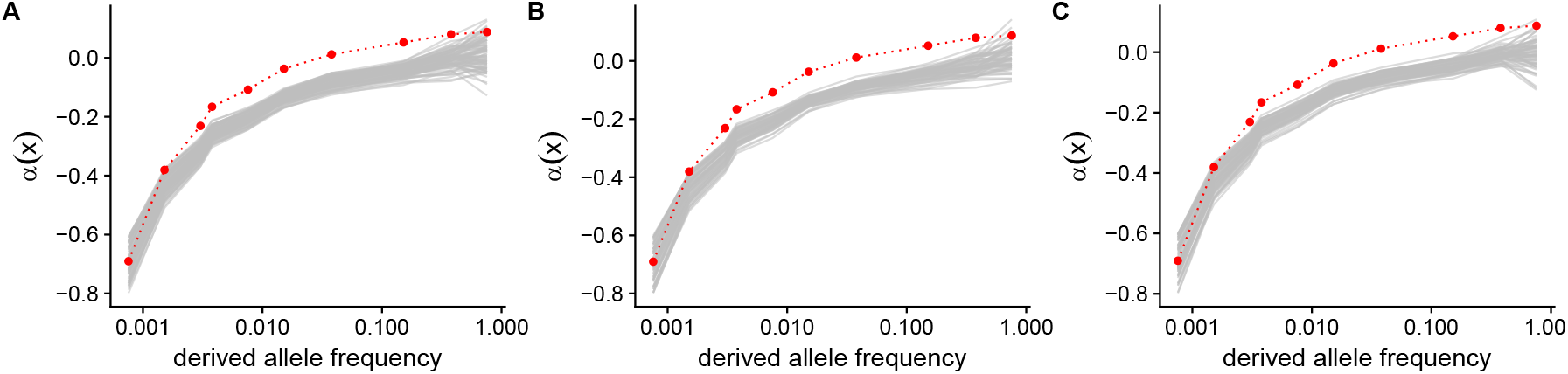
Summary statistics for simulations with very low adaptation (*α* < 0.01) as compared to the observed data for three different DFEs of alleles driving BGS. In A, alleles in flanking regions around genes have 2*Ns* = −10, in B 2*Ns* = −500, and in C 2*Ns* is a gamma-distributed mixture of weakly deleterious and strongly deleterious alleles (*37*).

**Figure S16:**
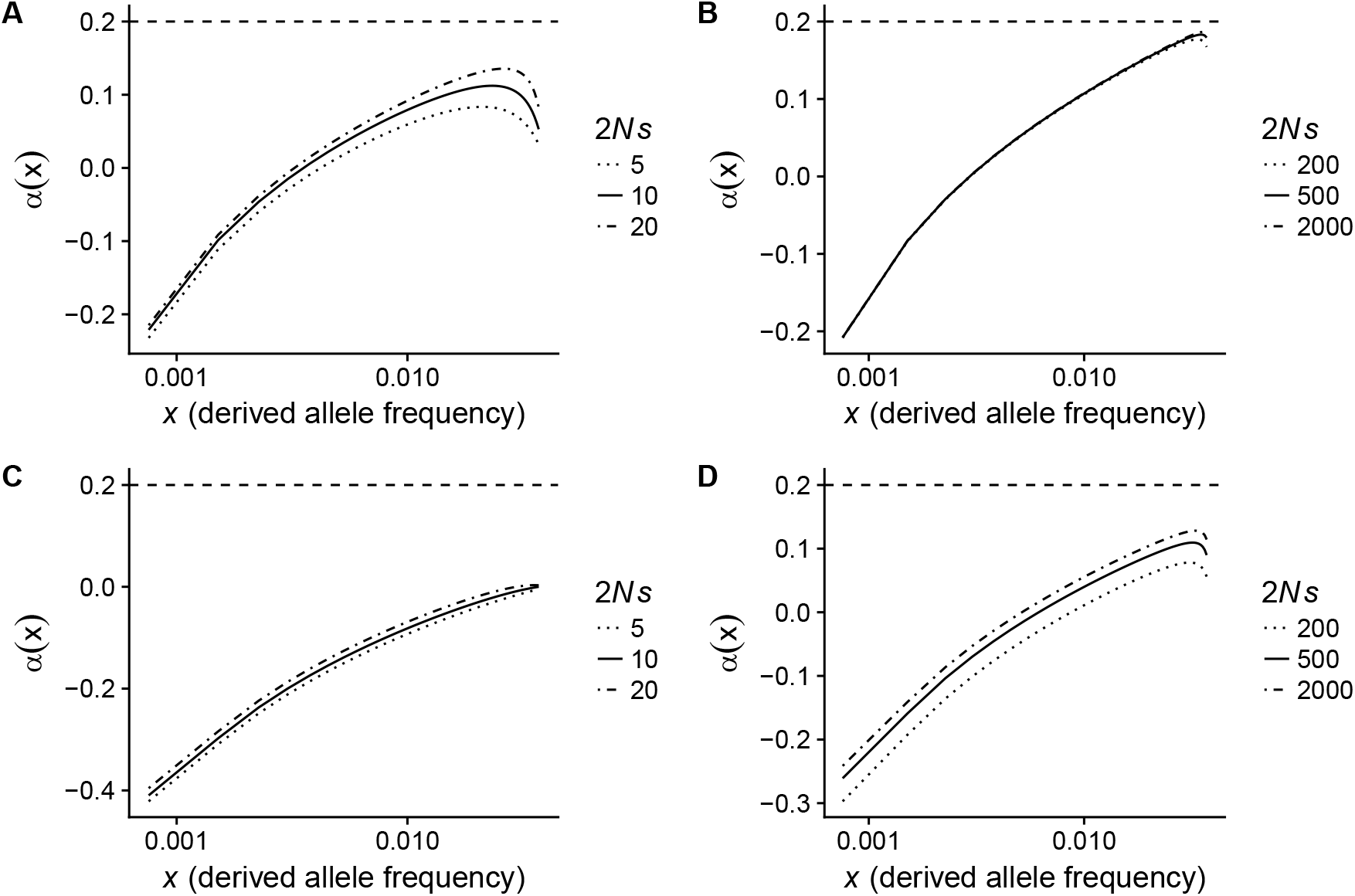
Comparison of analytical approximations to *α*(*x*) summary statistics for a range of 2*Ns* values at adaptive alleles. The dashed line at *α*(*x*) = 0.2 represents the true *α* in the absence of BGS. In A&B, there is no BGS, while 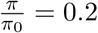 in C&D.

**Figure S17:**
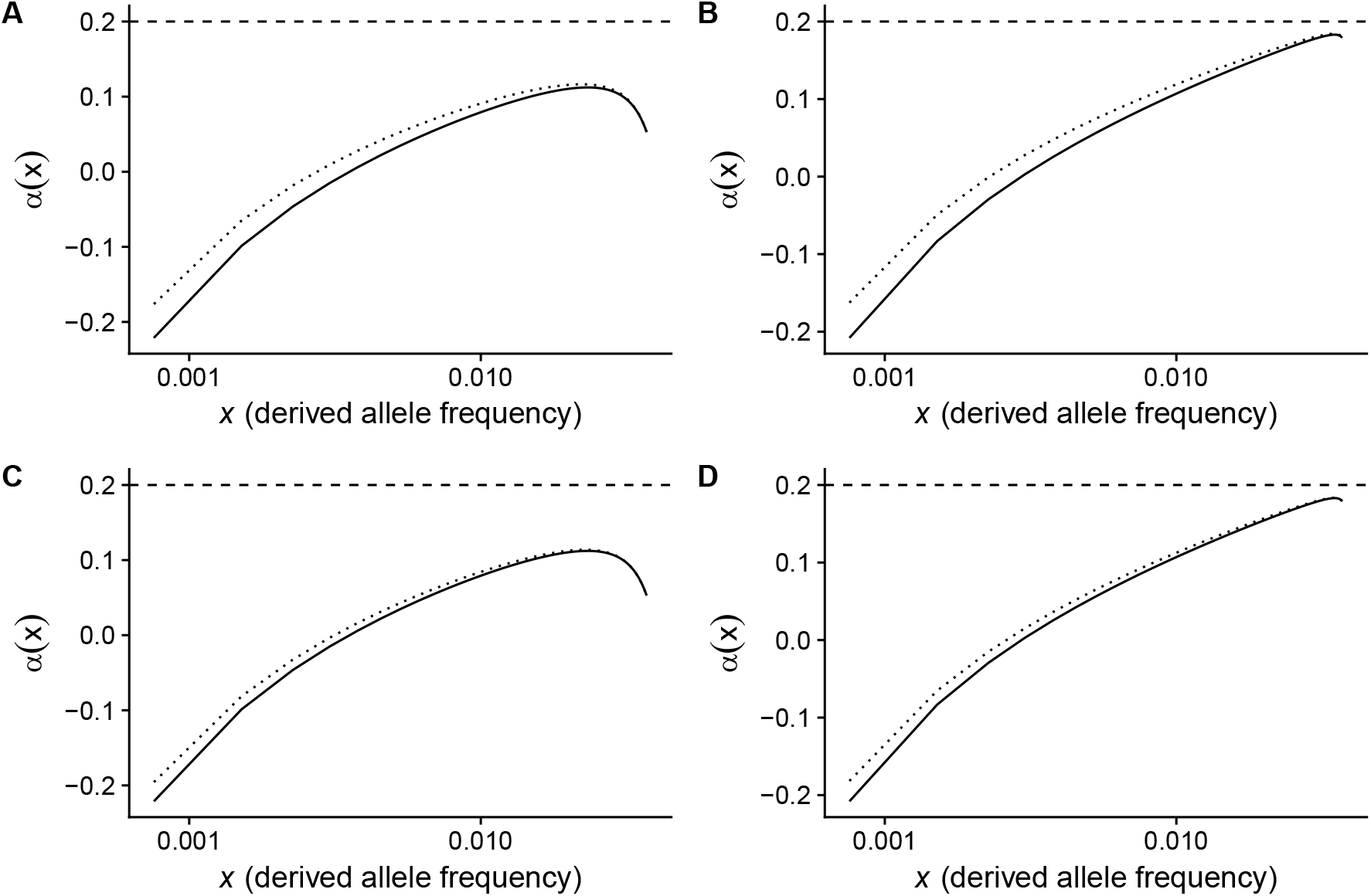
Analytical approximation to *α*(*x*) summary statistics when selection acts on synonymous alleles (dotted line) or does not act on synonymous alleles (solid line). In all panels, the true rate of adaptation *α*= 0.2 and 70% of synonymous alleles are assumed to be neutral. A&B: 29% of synonymous alleles are moderately deleterious and 1% are strongly deleterious. C&D 27% of synonymous alleles are moderately deleterious and 3% are strongly deleterious. In the left column, 2*Ns* = 10 for adaptive alleles, and in the right column 2*Ns* = 500 for adaptive alleles. These DFEs for synonymous alleles were motivated by Huang & Siepel (*62*), who estimated the DFE of synonymous alleles from human genomic data (*62*).

**Figure S18:**
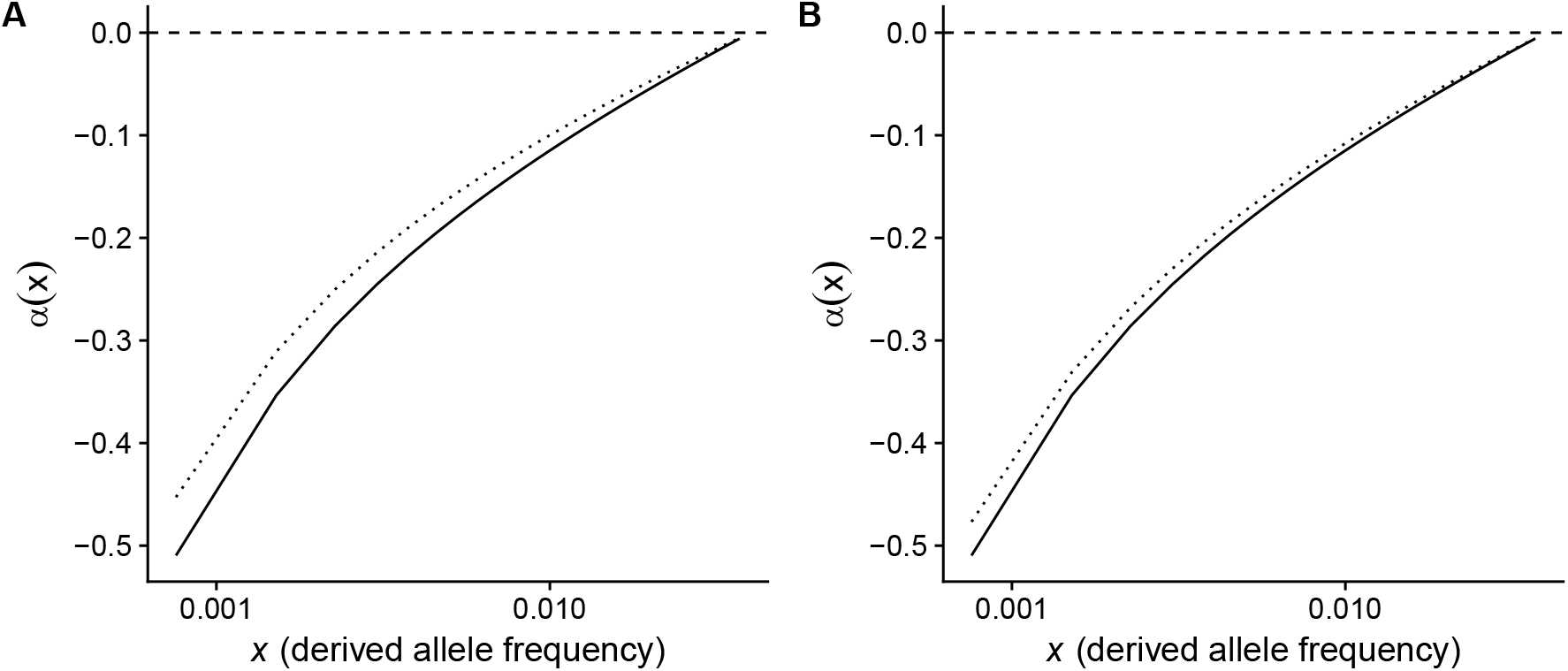
Analytical approximation to *α*(*x*) summary statistics when selection does (dotted line) or does not (solid line) act on synonymous alleles. In both panels, the true rate of adaptation *α* = 0.0 and 70% of synonymous alleles are assumed to be neutral. A: 29% of synonymous alleles are moderately deleterious and 1% are strongly deleterious. B 27% of synonymous alleles are moderately deleterious and 3% are strongly deleterious. These DFEs were motivated by an attempt to qualitatively match the distribution of Huang & Siepel, who estimated the DFE of synonymous alleles from human genomic data (*62*).

**Figure S19:**
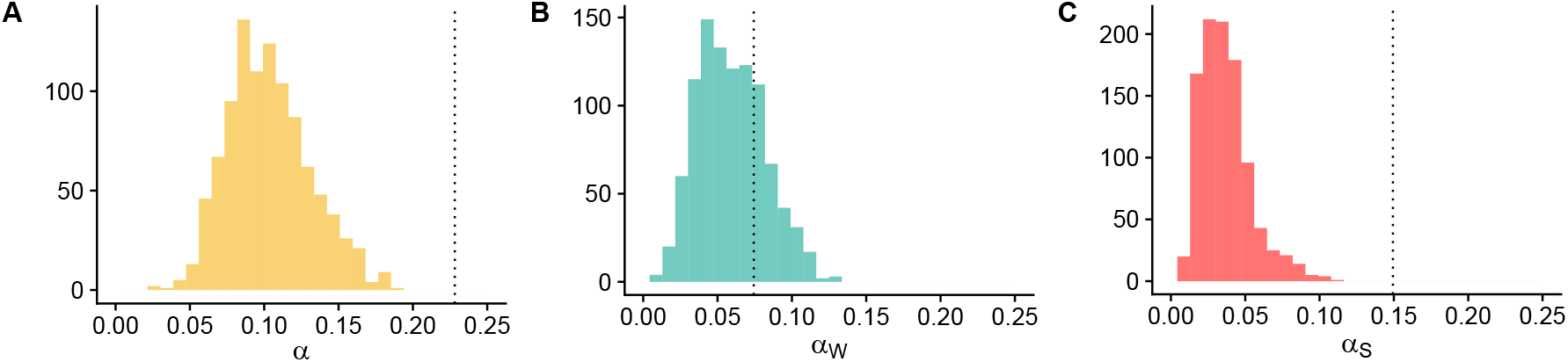
Median posterior estimates of *α*, *α_W_*, and *α_S_* for both VIPs (black dashed lines) and 1,000 independent sets of genes sampled from non-VIPs to approximately match VIPs for 13 potential confounding variables (histograms). For the set of VIPs that we identified, we built an equal-sized control set of non-VIPs that that has the same overall average values for 13 factors (DS, PN, PS, GC content, recombination rate, coding sequence length, average expression across 53 GTEx tissues, average GTEx expression in testis, average GTEx expression in lymphocytes, the number of protein-protein interactions, McVickers B for background selection, the coding sequence density, the overall density of PhastCons conserved elements). We randomly sampled non-VIPs such that each of the 13 factors does not depart on average by more than ±5% of their average for VIPs.

**Figure S20:**
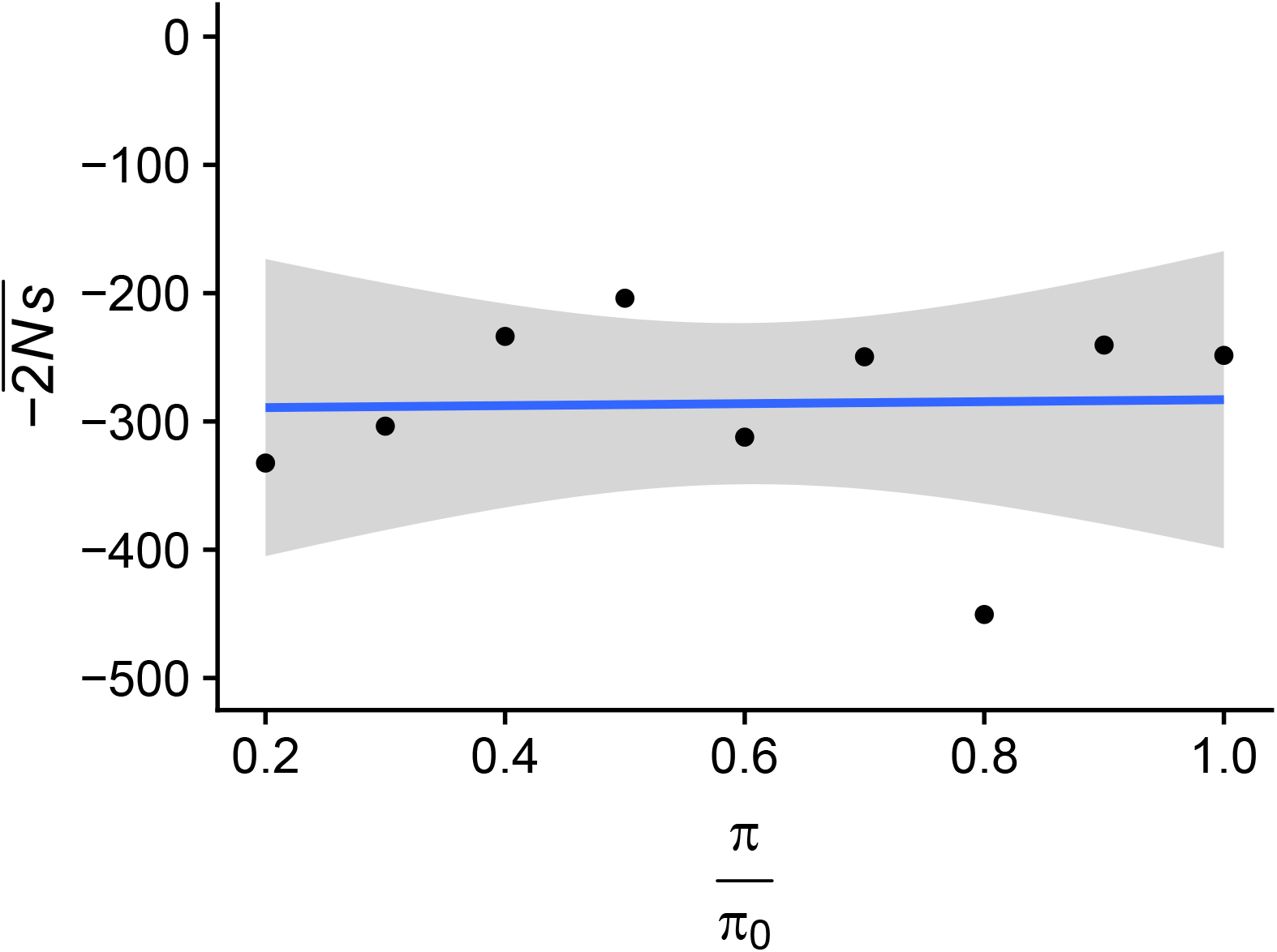
Posterior estimates of mean strength of negative selection 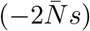 against deleterious nonsynonymous alleles as a function of estimated BGS strength. The line shows the best fit linear regression curve as determined by the method `geom_smooth` in the package `ggplot2`. The estimated correlation coefficient is *ρ* = 0.028.

